# Histone H4 acetyl-methyllysine marks accessible chromatin that resists compaction

**DOI:** 10.64898/2026.04.17.718779

**Authors:** Andreas P. Pintado-Urbanc, Charlie L. Brown, Leah J. Connor, Joshua T. Young, Yeil Kim, Elizabeth M. Black, Lilian Kabeche, Matthew D. Simon

**Author notes:** These authors contributed equally to this work (Charlie L. Brown, Leah J. Connor).

## Abstract

Certain regulatory DNA regions remain accessible even under conditions of widespread chromatin compaction. These regions are often marked by specific protein factors and histone modifications that help maintain their accessibility. Here, we examine the genomic landscape of acetyl-methyllysine (Kacme), a recently discovered histone post-translational modification. Across multiple systems, Kacme is highly enriched at sites of accessible chromatin, including active promoters, enhancers, silencers, and CTCF-binding sites. We find that Kacme is selectively retained at loci that resist condensation during mitosis, marks *XIST* and escapee regions on the inactive X chromosome in female cells and demarcates the boundaries of broad heterochromatin domains. Kacme-marked insulator elements block heterochromatin spreading and protect adjacent genes from transcriptional repression, even when H3K27me3 levels are pharmacologically elevated through KDM6A/6B inhibition. Taken together, our findings establish the chromatin features associated with Kacme and support a model in which Kacme helps safeguard chromatin accessibility at loci that resist compaction.

## Main Text

As cells adapt to their environment, grow, and differentiate, large regions of chromatin often undergo compaction. Even within condensed chromatin, however, certain genes and regulatory loci remain accessible. Specific histone modifications help maintain accessibility by opposing chromatin compaction through diverse mechanisms of action (*1–5*). For example, histone H4 lysine 16 acetylation (H4K16ac) counteracts chromatin compaction *in vitro* by weakening nucleosome-nucleosome interactions (*6, 7*), and exerts anti-silencing functions across yeast, fly, and human (*8–12*). Some histone post-translational modifications (PTMs) directly antagonize the deposition of repressive chromatin marks; specifically, H3K4 methylation can inhibit DNA methyltransferase activity (*13, 14*), and H3K36 methylation can block the polycomb repressive complex 2 (PRC2) from installing H3K27 methylation (*15–18*). Other histone PTMs, including H4K5ac and H3K27ac contribute to anti-silencing, for instance by marking genes during mitosis to facilitate rapid transcriptional reactivation in G1 (*19–22*). In addition to acetylation, the histone variant H2A.Z has been reported to maintain promoter accessibility and euchromatin integrity by restricting the spread of silent heterochromatin into active genomic regions (*23–25*). These examples highlight the pivotal roles of chromatin modifications in opposing compaction and motivate further investigation into the mechanisms and additional PTMs that sustain chromatin accessibility.

We recently reported the discovery of acetyl-methyllysine (Kacme) (*26*), a modification of histone H4 with regulatory properties distinct from those of methylation or acetylation alone, inspiring investigation for its physiological roles (*27, 28*). This work established Kacme as an abundant chromatin mark with levels comparable to those of activating PTMs such as H3K4me3, H3K27ac, and H4Kac, and involved the development of two highly validated antibody preparations: one recognizing Kacme across all tested peptide contexts, and another specific to Kacme within histone H4 tail peptides (H4Kacme). Using these reagents, we demonstrated that Kacme is primarily found at residues H4K5 and H4K12 and is enriched at the transcription start sites (TSSs) of thousands of actively transcribed genes. TSSs marked by Kacme are associated with increased rates of transcriptional initiation, and H4Kacme can be recognized by chromatin readers including the bromodomain-containing proteins BRD2 and BRD3.

While the chemistry of Kacme resembles that of neutral Kac, we found that H4K5ac, but not H4K5acme, can be removed by HDAC1 and HDAC3 *in vitro*, suggesting that Kacme may exhibit differential sensitivity to erasure relative to Kac. Furthermore, ChIP-seq studies in *D. melanogaster* S2 cells revealed that Kacme levels are differentially regulated relative to H4Kac under heat-stress conditions, indicating that Kacme likely encodes information distinct from acetylation alone. Indeed, recent studies in differentiating muscle cells showed that MYOD-mediated chromatin compaction was associated with a loss of Kacme but not H3K27ac at a subset of super-enhancers regions (*27*), suggesting a broader role for Kacme in regulating chromatin accessibility than previously appreciated.

In this study, we investigate the genomic landscape of Kacme relative to other chromatin marks in both steady-state and differentiating cellular models. We find that Kacme is anti-correlated with repressive modifications such as H3K27me3 and CpG methylation, and is strongly enriched at regions of open chromatin, including active promoters and enhancers targeted by pioneer transcription factors (TFs). During cellular differentiation, Kacme levels are dynamically reorganized, mirroring changes in chromatin accessibility and gene expression independently of H3 and H4 acetylation. These observations led us to hypothesize that Kacme functions to help maintain local chromatin accessibility–even under conditions of widespread compaction or transcriptional silencing. To test this hypothesis, we examined Kacme distribution (1) across mitosis and early G1, (2) at genes that escape X-chromosome inactivation (XCI), and (3) at heterochromatin boundary regions. We further demonstrate that Kacme-marked boundary elements limit H3K27me3 spreading when the H3K27me3 demethylases KDM6A and KDM6B are inhibited. Taken together, our findings support a model in which Kacme contributes to the maintenance of chromatin accessibility at regulatory sites, particularly in contexts associated with broad chromatin compaction and gene silencing.

### Kacme marks accessible promoters independent of transcription

Kacme is associated with active promoters and transcriptional initiation (*26*), suggesting a role in establishing or maintaining open chromatin upstream of RNA Polymerase II (Pol II) recruitment (*1, 29–33*). To investigate the relationship between Kacme-marked regions and chromatin accessibility, we performed transposase-accessible chromatin sequencing (ATAC-seq) in HEK293T cells and compared accessibility profiles with Kacme enrichment. Promoters with higher levels of Kacme exhibited increased accessibility (Fig. 1A-B), consistent with previous observations linking Kacme dynamics to changes in ATAC-seq signal during skeletal myogenesis (*27*). Kacme-enriched promoters were depleted of heterochromatin-associated marks such as H3K27me3 (Fig. 1C). Even after controlling for H4K5ac levels, higher Kacme levels remained strongly associated with increased ATAC-seq signal at the TSS (Fig. 1D). Similar analyses performed in an independent human cell line (monocytic THP-1 cells) support the generality of the association between promoter-proximal Kacme and chromatin accessibility (Fig. S1A-B).

**Fig. 1.**
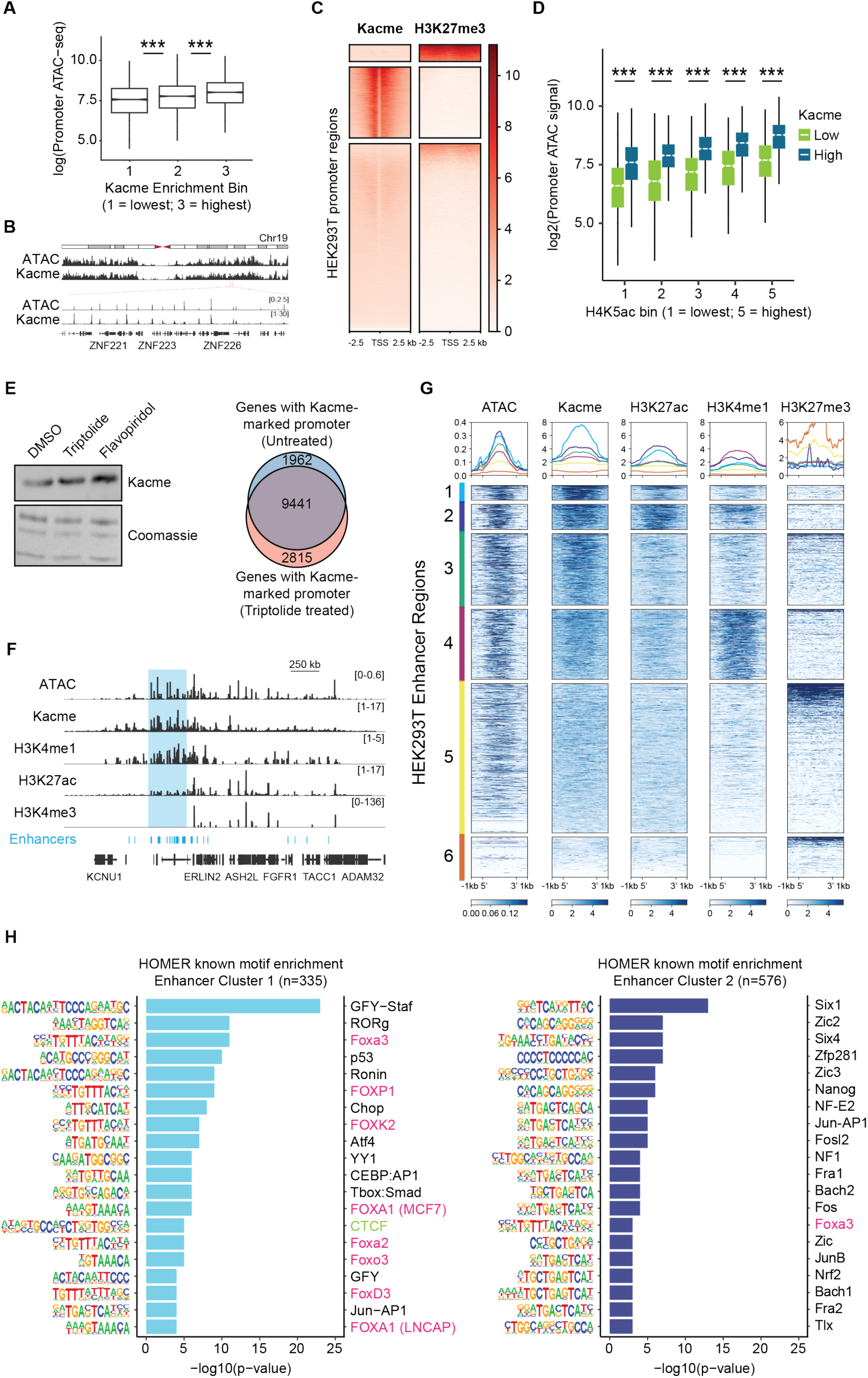
Kacme marks regions of accessible chromatin, including enhancers targeted by pioneer TFs. (**A**) Relative promoter accessibility (±50 bp of the TSS) in HEK293T cells as determined by ATAC-seq. Normalized ATAC-seq signal was binned by promoter Kacme levels (±1 kb of the TSS) as determined by ChIP-seq. (**B**) Genome browser visualization of ATAC-seq and Kacme ChIP-seq profiles at selected loci on chromosome 19 in HEK293T cells. (**C**) Heatmaps of Kacme ChIP-seq and H3K27me3 CUT&RUN signal centered on transcription start sites (±2.5 kb) across all annotated HEK293T promoter regions. (**D**) Relative promoter accessibility in HEK293T cells for genes in the 25^th^ or 75^th^ percentiles of Kacme CUT&RUN levels. Data are binned by H4K5ac CUT&RUN signal. (**E**) Left: Representative western blot of histones from HEK293T cells treated with the transcription inhibitors triptolide (10 μM, 1 h) or flavopiridol (500 nM, 1 h). Right: Venn diagram showing the overlap between genes with a promoter-proximal Kacme peak (±3 kb of the TSS) in untreated HEK293T cells (light blue) and in triptolide-treated HEK293T cells (light red). Kacme peaks were identified using MACS2. (**F**) Genome browser tracks showing ATAC-seq, Kacme, H3K4me1, H3K27ac, and H3K4me3 signal profiles across an enhancer-rich locus in HEK293T cells. (**G**) Heatmaps and aggregated signal profiles for ATAC-seq, Kacme, H3K27ac, H3K4me1, and H3K27me3 across six enhancer clusters identified in HEK293T cells by k-means clustering. Aggregated profiles show the mean signal across enhancer regions within each cluster. (**H**) HOMER known motif analysis for enhancer clusters 1 and 2 (Fig. 1G), showing the top 20 enriched motifs with their best-match transcription factors and associated p-values. FOX proteins highlighted in magenta, CTCF highlighted in green. (**A, D**) Distribution means compared with two-tailed unpaired Wilcoxon test for two biological replicates. *** = p < 0.001.

Our data are consistent with the hypothesis that Kacme functions upstream of transcriptional initiation; however, we also considered the alternative possibility that Kacme deposition occurs as a consequence of ongoing transcription (*34–36*). To test whether transcription is necessary for the association between Kacme and accessible chromatin, we inhibited transcription at two distinct steps of transcriptional initiation using the small molecule inhibitors triptolide and flavopiridol and monitored Kacme levels. We found that global Kacme levels in HEK293T cells were independent of downstream transcriptional activity (Fig. 1E) and that Kacme distribution was largely unchanged following acute transcriptional inhibition (Fig. 1E and Fig. S1C). Together, these results support the model that Kacme acts upstream of transcriptional activation at open promoters.

### Kacme marks additional chromatin regions, including accessible enhancers and silencers

The association between Kacme and chromatin accessibility is not limited to TSSs. We observe Kacme enrichment at numerous accessible regions distal to gene TSSs in both HEK293T and THP-1 cells (Fig. S2A-B). This observation is consistent with recent work showing that Kacme localizes to accessible MYOD-bound super-enhancers in IMR90 cells (*27*), supporting a model in which Kacme exerts regulatory functions beyond transcriptional initiation. Further supporting this model, analysis of Kacme distribution in HEK293T cells revealed that a substantial fraction (∼30%) of Kacme-enriched sites lie more than 1 kb from an annotated TSS, with enrichment observed in introns, coding exons, and distal intergenic regions (Fig. S2C). A similar pattern was observed in THP-1 cells (Fig. S2D), indicating that promoter-distal Kacme enrichment is a general feature across human cell types.

We compared Kacme-enriched regions in HEK293T cells to enhancer annotations obtained from the EnhancerAtlas database (*37*). We excluded enhancers with overlapping H3K4me3 peaks from our analyses to avoid cases where alternative TSSs were misannotated as enhancer regions (*32*). We found that Kacme was enriched at accessible enhancers in HEK293T cells (Fig. 1F and Fig. S2E), consistent with its localization to active enhancer ChromHMM states in K562 cells (Fig. S2F). Notably, Kacme showed differential enhancer distribution relative to canonical enhancer-associated marks such as H3K4me1 and H3K27ac (Fig. 1F and Fig. S2G). K-means clustering of ChIP-seq enrichment across all HEK293T enhancer regions identified a subset of highly accessible enhancers (Cluster 1, n = 335) preferentially marked by Kacme relative to H3K27ac and H3K4me1 (Fig. 1G). These enhancer regions showed strong motif enrichment for forkhead box (FOX) TFs (Fig. 1H), which have been implicated in pioneer activity and the establishment of accessible chromatin (*38*). Analysis of Kacme ChIP-seq in fly S2 cells (*26*) supports the evolutionary conservation of Kacme at enhancer regions (Fig. S2H-I). In addition to enhancer elements, we observed Kacme enrichment at a subset of annotated silencer regions that exhibit chromatin accessibility, indicating that Kacme can also mark accessible regulatory loci with repressive activity (Fig. S2J-K). We conclude that Kacme marks diverse regions of accessible chromatin, including TSSs, enhancers, super-enhancers, and silencers.

Given its widespread genomic distribution, we hypothesized that Kacme may define a chromatin signature distinct from that of other canonical activating histone marks. To explore this possibility, we compared our Kacme enrichment profile with publicly available ENCODE ChIP-seq datasets in human K562 cells. We included transcriptional machinery, TFs, and common histone PTMs, including H3K9ac, H3K27ac, H4Kac, H3K4me1, and H3K4me3 (*39, 40*). Using the chromatin state discovery tool ChromHMM (*41*), we segmented the K562 genome into a 15-state model, capturing a range of active and repressed chromatin environments (Fig. 2A). We found that one state (State 12, Fig. 2A) exhibited strong, preferential enrichment for Kacme, suggesting that certain genomic regions can be distinguished by the presence of Kacme. State 12, while preferentially marked by Kacme, also harbored other promoter- and enhancer-associated marks, along with anti-silencing marks (e.g. H3K27ac, H3K4me1, H4Kac, and H2A.Z). Kacme peaks associated with State 12 showed significant overlap with CpG islands (Fig. 2B), and subsequent integration of Kacme ChIP-seq data with publicly available reduced representation bisulfate sequencing (RRBS) datasets revealed a strong correlation with CpG hypomethylation (Fig. 2C). This relationship was independent of local acetylation levels (Fig. S2L-M), suggesting that regions preferentially marked by Kacme retain hallmarks of an active chromatin signature (*42, 43*).

**Fig. 2.**
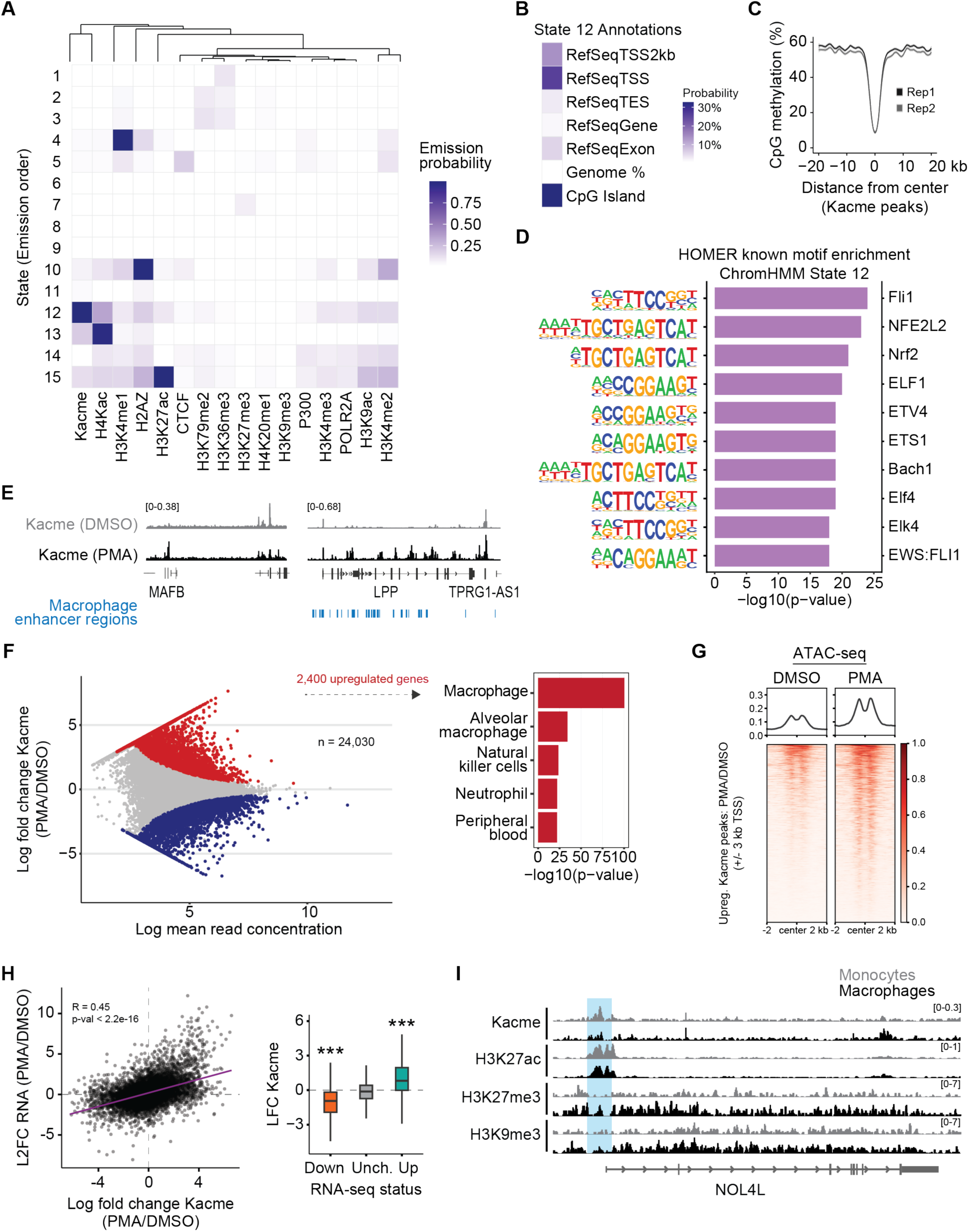
Kacme defines an accessible chromatin state linked to transcriptional activation, including during macrophage differentiation. (**A**) Histone mark and transcription factor emission probabilities, along with chromatin state definitions, for the 15-state ChromHMM model generated in wild-type K562 cells. Consensus peak BED files for histone PTMs and transcription factors in K562 cells were used as input for ChromHMM. Features were hierarchically clustered using Euclidean distance and complete-linkage clustering, while chromatin states were displayed in numerical order. (**B**) Genomic annotation enrichment for ChromHMM State 12, showing relative overlap with gene features and CpG islands. (**C**) Metaplot analysis of average CpG methylation levels centered on consensus Kacme peak summits in HEK293T cells. LOESS-smoothed curves with 95% confidence intervals show the mean signal ± CI across 100-bp bins spanning ± 20kb around the peak center for each biological replicate. (**D**) HOMER known motif analysis for ChromHMM State 12 genomic regions. The top 10 enriched motifs are ranked by significance with their best-match transcription factors and associated p-values. (**E**) Genome browser tracks showing Kacme ChIP-seq signal at the *MAFB* and *LPP* loci in THP-1 cells treated with DMSO or PMA. Blue bars denote macrophage-specific enhancer regions, as defined by EnhancerAtlas 2.0. (**F**) Left: MA plot depicting the changes in promoter Kacme ChIP-seq signal in THP-1 cells following PMA treatment. Promoters with significantly differential Kacme levels are indicated in red (L2FC > 0, FDR-adjusted p < 0.05) and blue (L2FC < 0, FDR-adjusted p < 0.05). Right: ARCHS4 Tissues enrichment analysis for genes with significantly upregulated promoter Kacme signal (n = 2,400 genes). (**G**) Normalized ATAC-seq signal changes following PMA treatment in THP-1 cells, plotted over significantly upregulated Kacme peaks (PMA/DMSO) located within 3 kb of gene TSSs. (**H**) Left: Scatterplot comparing the changes in promoter Kacme enrichment and RNA-seq levels following PMA treatment in THP-1 cells. Pearson correlation analysis was performed to assess the changes in L2FC values across experimental datasets. Right: Boxplots comparing changes in promoter Kacme levels at genes downregulated (L2FC < −1, FDR-adjusted p < 0.05), unchanged, or upregulated (L2FC > 1, FDR-adjusted p < 0.05) by RNA-seq following PMA treatment. Statistical significance between groups was determined using pairwise Wilcoxon rank-sum tests. *** = p < 0.001. (**I**) Genome browser tracks of Kacme, H3K27ac, H3K27me3, and H3K9me3 ChIP-seq profiles in THP-1 monocytes versus macrophages at the *NOL4L* locus.

TF motif enrichment on regions associated with the ChromHMM State 12 output revealed significant enrichment for TFs belonging to the ETS family, including FLI1, ELF1, ETV4, ETS1, and ELF4 (Fig. 2D). ETS proteins have been linked to diverse biological processes (*44*) and have been reported to act as pioneer transcription factors (*45*). Together, these results suggest that Kacme serves as a principal chromatin signature for a subset of accessible regulatory regions, where it may facilitate transcription factor binding and modulate downstream gene expression.

### Kacme redistribution during THP-1 differentiation reflects activation of lineage-specific programs

We next sought to understand how Kacme distribution relates to chromatin accessibility across cell fate transitions. Previous work has shown that Kacme is associated with the activation of gene expression pathways linked to cell differentiation (*27*). To investigate this further, we leveraged the widely used THP-1 monocytic differentiation model (*46, 47*). Treatment of leukemic THP-1 cells with phorbol 12-myristate-13-acetate (PMA) induces monocyte-macrophage differentiation, characterized by widespread changes in histone PTM profiles, chromatin accessibility, and transcription (*47–49*), providing a framework to examine the distribution of Kacme under dynamic conditions.

As expected, PMA treatment of THP-1 cells for 72 hours resulted in characteristic morphological changes (Fig. S3A) and robust transcriptional activation of macrophage-associated genes such as *DLX3*, *EGR3*, and *MAFB* (Fig. S3B). ChIP-seq profiling revealed that Kacme localization is extensively rewired following PMA-induced differentiation (Fig. 2E-F), which we confirmed by ChIP followed by quantitative PCR (ChIP-qPCR) (Fig. S3C). We also validated these results through ChIP-seq experiments using an H4Kacme-specific antibody (*26*), observing strong concordance with the results obtained using the pan-Kacme antibody (Fig. S3D).

We identified 2,400 genomic loci with significantly increased promoter Kacme levels after differentiation (Fig. 2F). The corresponding genes were primarily associated with macrophage-relevant processes such as neutrophil degranulation, cellular import, and actin cytoskeleton remodeling (Fig. 2F and Fig. S3E) (*50*). We observed evidence of *de novo* Kacme installation at TSSs and enhancer regulatory domains (Fig. 2E), with striking gains in Kacme levels at the promoters of important macrophage transcriptional regulators (Fig. S3F) (*51*). Conversely, analysis of the 2,824 loci that significantly lost Kacme signal revealed a strong enrichment for cell cycle and DNA metabolism related processes (Fig. S3G). This can be observed at genes such as *GFI1*, *MYB*, and *IRF8* (Fig. S3H), which all encode factors involved in the maintenance of immature myeloid proliferation (*52–54*).

To systematically assess the relationship between Kacme, accessibility, and gene expression, we integrated matched ATAC-seq and RNA-seq datasets from PMA-treated THP-1 cells (*49, 55*). Genes exhibiting increased Kacme levels at promoters showed enhanced ATAC signal (Fig. 2G) and robust transcriptional upregulation (Fig. 2H), indicating a strong correlation between differentiation-induced gains in Kacme and chromatin activation. We next compared Kacme localization changes to those of H3K27ac, a mark previously implicated in the activation of promoters and enhancers during THP-1 differentiation (*49*). While PMA-induced changes in Kacme and H3K27ac are largely correlated, differential binding analysis revealed subsets of genes with distinct regulation of Kacme and H3K27ac (Fig. S4A-B). Notably, genes that gain Kacme but not H3K27ac (n = 548 genes) show marked transcriptional upregulation (Fig. S4C) and enrichment for macrophage-related transcription factors and biological pathways (Fig. S4D). In contrast, genes gaining H3K27ac but not Kacme (n = 115 genes) do not display such an enrichment (Fig. S4E). Finally, genes that lose Kacme but retain H3K27ac (n = 342 genes) are generally downregulated (Fig. S4F). These observations suggest that Kacme contributes to transcriptional activation during monocyte-macrophage differentiation independently of other canonical activating acetylation marks. Consistently, H4K5ac ChIP-seq experiments performed under matched conditions (+/- PMA) also revealed differences between changes in Kacme and H4K5ac (Fig. S4G). Taken together, these data indicate that Kacme is regulated in concert with changing transcriptional programs and may promote or reinforce active chromatin states essential for myeloid differentiation.

Finally, we examined the relationship between Kacme and repressive histone modifications during THP-1 differentiation. Changes in Kacme levels correlated with altered patterns of heterochromatin organization (Fig. 2I and Fig. S4H-I). For example, at the *NOL4L* locus, PMA-induced loss of Kacme coincided with gains of H3K27me3 and H3K9me3 across the promoter and gene body (Fig. 2I). Collectively, our analyses of Kacme across multiple cell types suggest that the presence of Kacme at accessible TSSs may reflect a more general function in which Kacme acts to oppose the formation of condensed chromatin.

This prompted us to examine the distribution and dynamics of Kacme at regions known to resist large-scale chromatin compaction, including (1) genes that undergo rapid transcriptional reactivation after mitosis, (2) genes expressed from the otherwise inactive X chromosome in female mammals, and (3) loci that resist H3K27me3 spreading at heterochromatin boundaries.

### Kacme marks accessible chromatin during the mitosis-G1 phase transition

During mitosis, chromatin is compacted and transcription is largely repressed (*56, 57*), yet some TFs remain bound on chromosomes (*58*). Certain H4 modifications, including H4K5ac, are retained at select genes, where they are thought to promote open chromatin conformations and facilitate rapid transcriptional reactivation upon entry into early G1 (*19, 20*). To test whether Kacme marks loci that remain accessible during mitosis, we first performed immunofluorescence studies against Kacme in chromosome spreads prepared from human cells arrested in nocodazole (Fig. 3A). We observed broad Kacme staining across chromosome arms, indicating that Kacme is at least partially retained during mitosis.

**Fig. 3.**
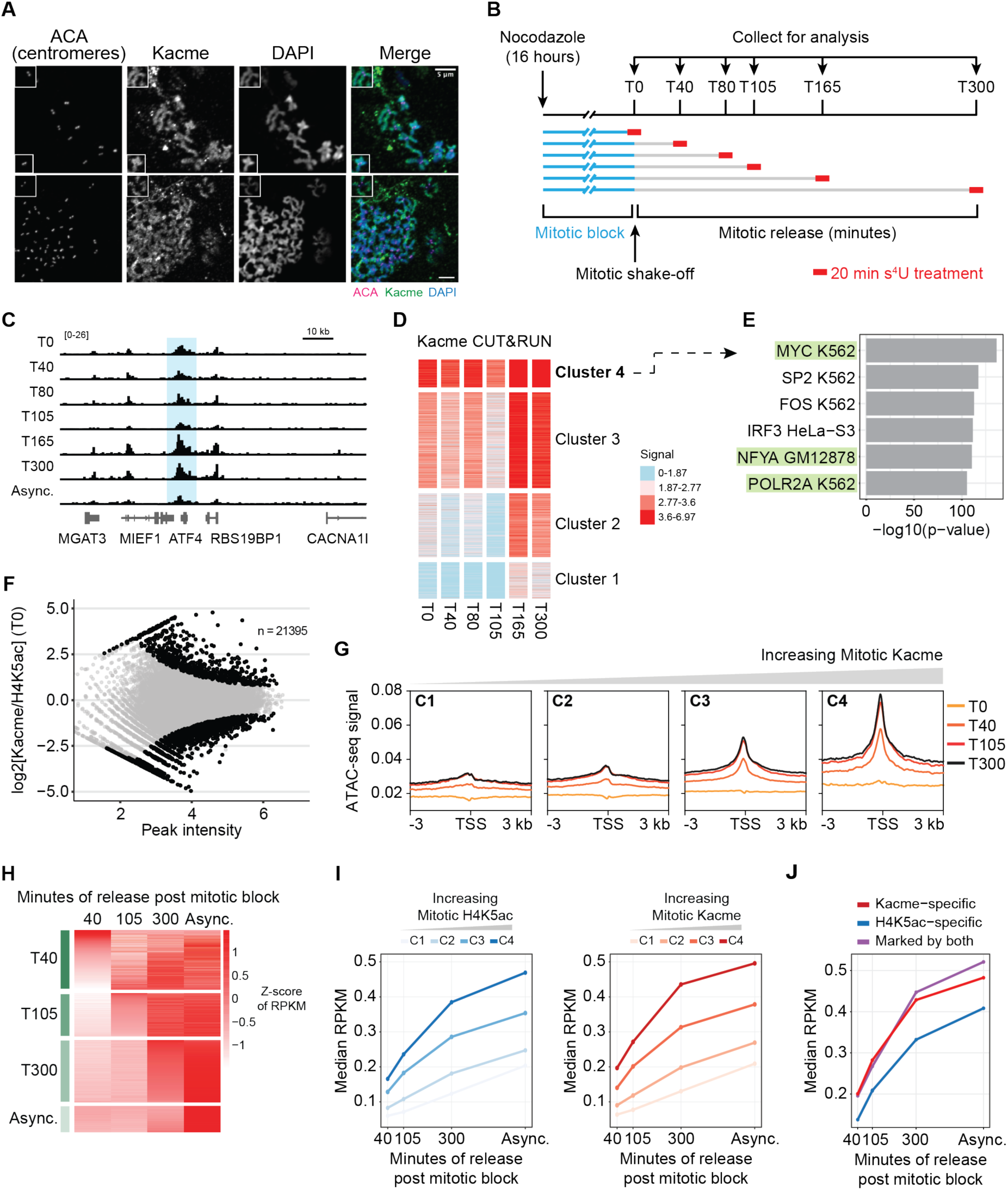
Kacme marks genes poised for rapid transcriptional reactivation at the mitosis-G1 phase transition. (**A**) Chromosome spreads were prepared from HeLa cells arrested in nocodazole and immunostained with antibodies against Kacme (green) and centromeres using anti-centromere antibody (ACA, magenta). DNA was counterstained with DAPI (blue). Insets show representative chromosome regions with Kacme retained on condensed mitotic chromatin. Scale bar, 5 μm. (**B**) Protocol to synchronize HEK293T cells and arrest at the G2/M boundary using a nocodazole block/mitotic shake-off. Matched CUT&RUN (against Kacme and H4K5ac), ATAC-seq, and nascent RNA sequencing were performed at several timepoints across the M/G1 transition. (**C**) Genome browser tracks showing a representative gene, *ATF4*, that is marked by Kacme throughout the mitosis-G1 phase transition. (**D**) Heatmap showing hierarchical k-means clustering of normalized Kacme CUT&RUN signal across promoter regions during mitotic release. Four major clusters (C1-C4) were identified, with 1,579 genes in C4 displaying the strongest Kacme signal during mitosis. Increasing signal intensity is indicated by a red color scale. (**E**) Transcription factor enrichment analysis of genes associated with Cluster 4 (Fig. 3D). Top enriched motifs include mitotically retained TFs such as MYC, NFYA, and POLR2A. (**F**) Differential binding analysis of regions of enrichment from Kacme and H4K5ac CUT&RUN in mitotically arrested cells (T0). Black points are regions of significant difference (p-value < 0.05). (**G**) Normalized ATAC-seq signal at the transcription start sites of genes associated with each Kacme cluster (C1-C4; Fig. 3D) across selected mitotic release time points (T0, T40, T105, T300). Promoters with the highest level of mitotic Kacme enrichment showed the most significant gain in accessibility upon release from mitosis into early-G1. (**H**) Heatmap showing nascent transcription levels (TimeLapse-seq) for genes that first reach ≥0.5-fold above asynchronous levels during mitotic release. Within each timepoint (columns), genes are rank-ordered based on their transcription levels. Nascent transcription values are shown as Z-scores of RPKM. (**I**) Right: Median RPKM of nascent transcription across mitotic release for genes in each Kacme cluster (C1-C4; Fig. 3D). Left: Median RPKM of nascent transcription across mitotic release for genes grouped by mitotic H4K5ac levels (C1-C4). (**J**) Comparison of nascent transcriptional dynamics at genes preferentially marked by Kacme, H4K5ac, or both, during the mitotic release time course. Median RPKM values of nascent transcription for genes associated with each set are shown across mitotic release timepoints.

To characterize the genome-wide localization of Kacme across the mitosis-G1 transition, we performed a mitotic release time course experiment in HEK293T cells (Fig. 3B). Nocodazole-mediated arrest, mitotic shake-off enrichment, and release into G1 were performed as previously described (*59*). The extent of mitotic arrest and reentry into G1 was evaluated by Western blot analysis of H3Ser10 phosphorylation (Fig. S5A) and by propidium iodide staining (Fig. S5B). We confirmed that Kacme, like H4K5ac, is maintained on chromatin throughout the mitosis-G1 phase transition (Fig. S5A). Comparative CUT&RUN experiments targeting Kacme and H4K5ac (Fig. 3B) revealed that Kacme is retained at a subset of TSSs (Fig. 3C) and enhancer regions (Fig. S5C-D) throughout the mitosis-G1 transition. Profiling of Kacme signal over promoter regions identified subsets of genes with differential enrichment in asynchronous and mitotically arrested cells (Fig. S5E), consistent with locus-specific preservation of Kacme during mitosis.

We defined four gene clusters based on promoter Kacme enrichment across the mitosis-G1 transition (Fig. 3D). Transcription factor enrichment analysis on the set of 1,579 genes with the highest Kacme signal (Cluster 4) revealed strong enrichment for binding by known mitotically retained TFs such as MYC, NFYA, and POLR2A (Fig. 3E) (*60–62*). We also observed mitotic Kacme localization at CTCF binding motifs (Fig. S5F-G), which may relate to CTCF’s reported retention at select loci during mitosis (*63*). Together, these results suggest that Kacme preferentially marks regions of mitotic chromatin that remain permissive to TF binding, supporting our central hypothesis.

We next compared Kacme and H4K5ac levels across the mitosis-to-G1 transition and identified subsets of promoters exhibiting significant relative enrichment differences (Fig. 3F). Genes with high relative Kacme-to-H4K5ac ratios during mitosis were enriched for DNA replication and repair pathways (Fig. S5H), as exemplified by the *Hist1* locus (Fig. S5I). Interestingly, we observed a global increase in Kacme levels at TSSs and enhancer regions upon exit from mitosis and entry into early G1, a trend not observed for H4K5ac (Fig. S5J-K). This increase may reflect a role for Kacme in re-establishing active chromatin states during the transcriptional spike that occurs upon G1 entry (*64*). Moreover, these results demonstrate that Kacme is dynamically regulated across the mitosis-G1 transition in a manner distinct from H4K5ac.

To investigate the relationship between Kacme and local chromatin architecture during the mitosis-G1 transition, we performed ATAC-seq at four representative timepoints spanning the mitotic release time course (0-, 40-, 105-, and 300-minutes post-arrest, similar to (*59*)). Global ATAC-seq levels were low in nocodazole-arrested cells, consistent with mitotic chromatin compaction, and gradually increased upon exit from mitosis (Fig. S6A). Despite the overall condensation of chromatin during mitosis, several studies have shown that specific gene regulatory elements retain local accessibility (*65*). Upon integration with the previously described CUT&RUN results (Fig. 3D), we found that mitotic Kacme levels were strongly predictive of TSS accessibility in nocodazole-arrested cells (Fig. S6B). Notably, genes with the highest mitotic Kacme enrichment (Cluster 4, n = 1,579) also displayed the most significant gain in promoter accessibility upon exit from mitosis and entry into early G1 (Fig. 3G). Genes with little to no mitotic Kacme (Cluster 1, n = 2,077) displayed limited accessibility changes throughout the 300-minute time course.

To examine transcriptional changes during the mitosis-G1 transition, we performed TimeLapse-seq (*66*) with short exposures to s^4^U (20 min) to capture changes in newly synthesized RNAs (Fig. 3B). Consistent with previous reports (*59*), gene expression was reactivated in distinct waves following exit from mitosis (Fig. 3H and Fig. S6C). Our results further recapitulate the finding that H4K5ac is associated with rapid transcriptional reactivation upon entry into G1 (Fig. 3I, (*19*)). Notably, increasing mitotic Kacme promoter signal was also associated with rapid transcriptional reactivation upon entry into G1 (Fig. 3I and Fig. S6D), independent of H4K5ac levels (Fig. 3J and Fig. S6E). Of the 505 genes with high relative mitotic Kacme-to-H4K5ac ratios (Fig. 3J), nearly half (43.8%) exhibited early reactivation within 40 minutes of mitotic release. In contrast, only 27.5% of genes with high relative H4K5ac-to-Kacme ratios were reactivated within the same time frame. Genes marked by Kacme that were reactivated early were strongly enriched for metabolic and translation-related pathways (Fig. S6F), whereas Kacme-marked genes with late reactivation (300 min) were associated with DNA replication and DNA damage response (Fig. S6G), indicating that Kacme may also serve as a long-term marker for genes involved in pathways relevant to the G1/S-phase transition (*22*).

These data support our model that Kacme contributes to the regulation of gene expression across the mitosis-G1 transition by maintaining promoter accessibility during mitosis. Genes marked by Kacme are primed for robust transcriptional reactivation upon entry into G1, suggesting that Kacme modifications represent a non-redundant means for cells to preserve regions of open chromatin in the face of widespread mitotic compaction.

### Kacme marks escapee genes on the inactive X chromosome (Xi)

We next examined the genomic distribution of Kacme across the X chromosomes in female cells subject to X-chromosome inactivation. XCI is the process by which one of the two X chromosomes in female mammals is randomly silenced to achieve dosage compensation (*67*). XCI involves coating of the inactive X chromosome (Xi) by the long non-coding RNA *XIST* and the deposition of silencing marks such as H3K27me3, H2AK119ub1, and DNA methylation (*68–71*). Despite the overall compaction of the Xi, a subset of genes, referred to as escapees, resist repression and remain transcriptionally active (*72, 73*). Inspection of bulk Kacme ChIP-seq data from female HEK293T cells revealed striking enrichment over the promoter and gene body of *XIST* (Fig. 4A), as well as X chromosome escapee genes such as *DDX3X, TBL1X,* and *KDM5C* (Fig. 4B). Kacme displayed preferential localization to the 5’ end of the *XIST* gene relative to other regions, especially when compared with canonical anti-silencing marks such as H3K27ac and H4Kac (Fig. 4A). Notably, we did not observe similar Kacme enrichment over *XIST* in THP-1 cells (Fig. 4C), likely reflecting the absence of XCI in this male-derived cell line.

**Fig. 4.**
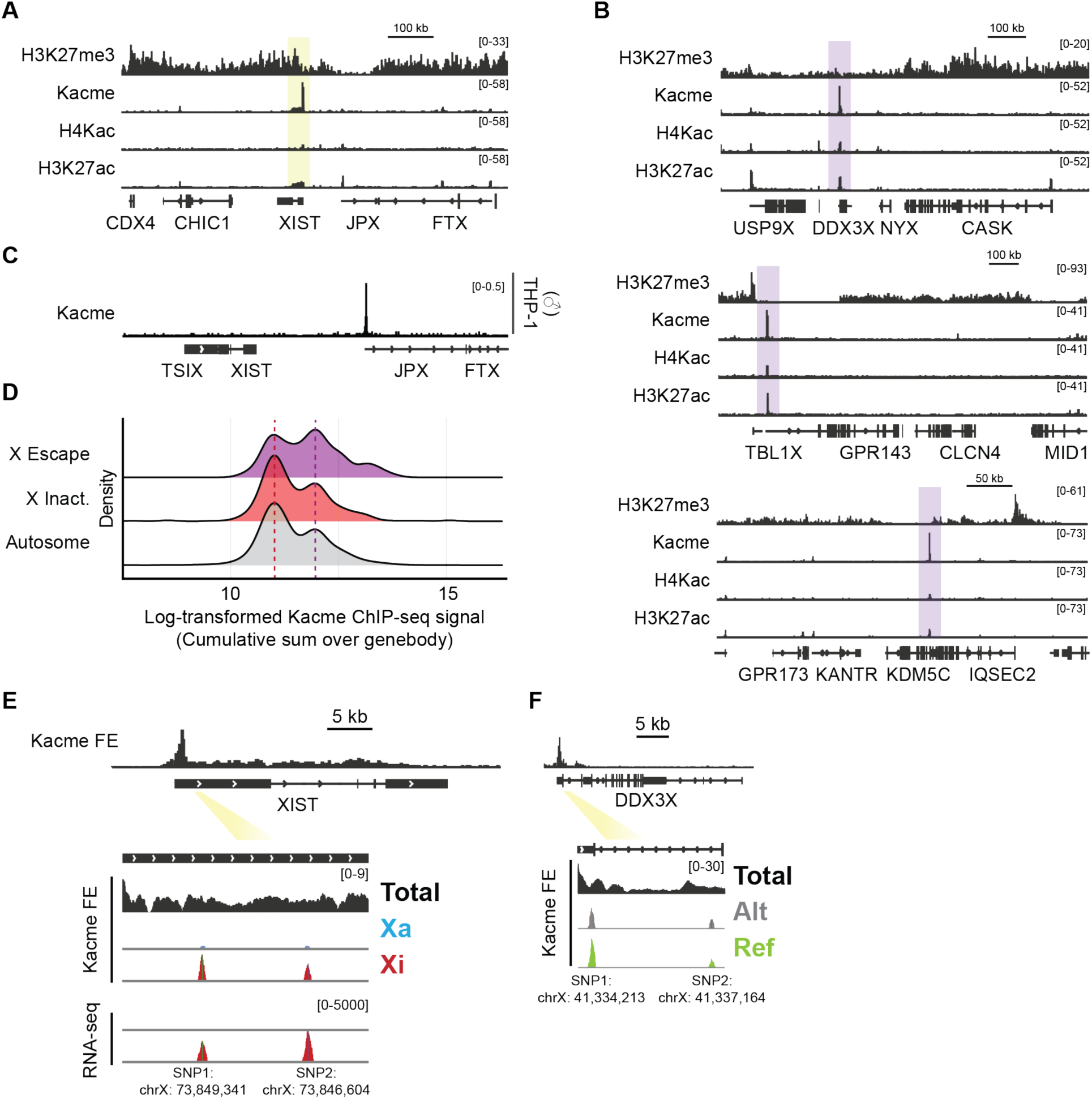
Kacme is enriched over *XIST* and escapee genes on the Xi chromosome in female cells. (**A**) Genome browser visualization of bulk H3K27me3, Kacme, H4Kac, and H3K27ac profiles at the X chromosome *XIST* locus in female-derived HEK293T cells. (**B**) Genome browser visualization of bulk H3K27me3, Kacme, H4Kac, and H3K27ac profiles at the escapee genes *DDX3X*, *TBL1X*, and *KDM5C* in female-derived HEK293T cells. (**C**) Genome browser visualization of bulk Kacme ChIP-seq signal at the *XIST* locus in male-derived THP-1 cells. (**D**) Distribution of log-transformed cumulative Kacme ChIP-seq signal across the gene bodies of X escapee, X inactivated, and autosomal genes. Dashed lines represent the maxima of the distribution for each category of genes. (**E**) Top: Bulk Kacme ChIP-seq enrichment across the *XIST* locus in untreated HEK293T cells. Bottom: Allelic contribution of Kacme ChIP-seq and RNA-seq reads mapping to the Xa (active X, blue) and Xi (inactive X, red) at *XIST* SNP1 and SNP2 positions. (**F**) Top: Bulk Kacme ChIP-seq enrichment across the *DDX3X* locus in untreated HEK293T cells. Bottom: Allelic contribution of Kacme ChIP-seq reads mapping to the reference (green) and alternative (grey) alleles at two *DDX3X* SNPs.

To determine whether Kacme is a characteristic feature of escapee loci in HEK293Ts, we examined Kacme ChIP-seq signal across constitutive escapee genes (*74*). We found that Kacme is significantly enriched at escapee genes compared to silenced X-linked genes (Fig. 4D). Taken together, these results support the notion that Kacme regulates regional resistance to heterochromatin spreading on the inactive X. To verify that Kacme marks *XIST* on the Xi, we leveraged heterozygous SNPs identified in HEK293T cells (*75*) to perform allele-specific analysis of Kacme ChIP-seq data at the *XIST* locus. Because *XIST* is expressed exclusively from the Xi in female somatic cells (*76, 77*), this analysis confirmed that Kacme reads mapping to the *XIST* gene body originate predominantly from the inactive X chromosome (Fig. 4E). In contrast, analysis of the escapee gene *DDX3X* revealed a more balanced allelic Kacme distribution (Fig. 4F), consistent with activation on both the active and inactive X chromosomes.

### Kacme localizes to CTCF binding sites and a subset of H3K27me3 domain boundaries

Consistent with previous observations, Kacme enrichment on the X chromosomes is anti-correlated with that of H3K27me3 (Fig. 4A-B). Notably, we found that Kacme marks the borders of numerous X chromosome H3K27me3 heterochromatin domains, even in the relative absence of other canonical anti-silencing modifications such as H3K27ac and H4Kac (Fig. S7A). More generally, genome-wide analysis of Kacme ChIP-seq signal at the edges of 4,162 broad H3K27me3 domains revealed a pronounced and localized enrichment peak (Fig. S7B), a finding which we validated by CUT&RUN against Kacme (Fig. 5A). Certain Kacme-enriched heterochromatin-euchromatin transitions lacked detectable H3K27ac (Fig. 5B), and we observed an overall preferential enrichment of Kacme relative to H3K27ac at these heterochromatin boundary regions (Fig. S7C-D).

**Fig. 5.**
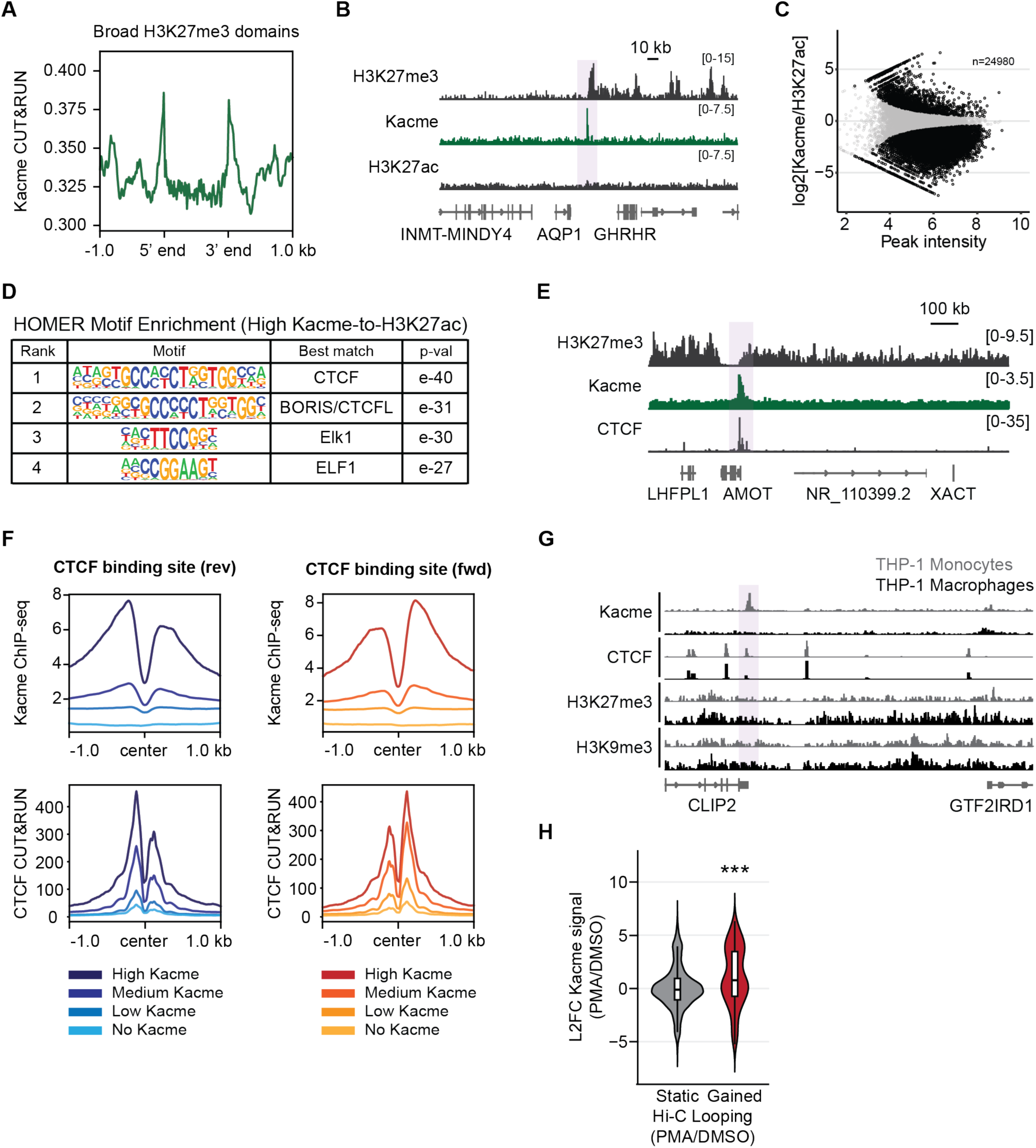
Kacme localizes to heterochromatin boundaries and CTCF-binding sites. (**A**) Metagene analysis of HEK293T Kacme CUT&RUN signal across broad H3K27me3 domains showing enrichment near domain boundaries. (**B**) Genome browser view of an example locus in HEK293T cells with Kacme enrichment at the border of a broad H3K27me3 heterochromatin domain. (**C**) Differential binding analysis of regions of enrichment from Kacme and H3K27ac ChIP-seq in untreated HEK293T cells. Black points are regions of significant difference (FDR-adjusted p < 0.05). (**D**) HOMER known motif analysis for regions with significantly high relative Kacme-to-H3K27ac ratios (Fig. 5C). The top two enriched motifs correspond to CTCF and CTCFL. (**E**) Genome browser view of the *AMOT* locus in HEK293T cells, which shows co-enrichment for Kacme and CTCF at the border of a broad H3K27me3 heterochromatin domain. (**F**) Metaplots of Kacme ChIP-seq and CTCF CUT&RUN signal centered on forward- and reverse-oriented FIMO-predicted CTCF binding sites (± 1kb). Sites are stratified by Kacme enrichment levels. (**G**) Genome tracks from DMSO- and PMA-treated THP-1 cells showing differences in Kacme, CTCF, H3K27me3, and H3K9me3 profiles at the *CLIP2* locus. The highlighted region marks a site that loses both Kacme and CTCF binding upon PMA-induced differentiation and exhibits spreading of heterochromatin. (**H**) Violin plots illustrating the L2FC in promoter Kacme ChIP-seq signal at genes that either remain static or exhibit significant increases in Hi-C chromatin looping upon PMA-induced differentiation of THP-1 cells (as reported in (*49*)). Statistical significance was determined using pairwise Wilcoxon rank-sum tests. *** = p < 0.001.

Our comparisons of Kacme with H3K27ac implicate Kacme as a mark enriched at CTCF binding sites. CTCF has been extensively studied for its role as an insulator that demarcates the boundaries of H3K27me3 heterochromatin domains (*78–80*). Moreover, while Kacme localization is generally highly correlated with H3K27ac, profiling of genomic regions with high relative Kacme-to-H3K27ac ratios (Fig. 5C) revealed a strong enrichment for CTCF- and CTCFL-binding motifs (Fig. 5D), supporting a role for Kacme in insulator function.

To investigate this relationship, we examined the distribution of Kacme across predicted CTCF-binding motifs in the human genome (n = 21,671 sites). We confirmed that Kacme is highly enriched over CTCF motif regions in HEK293T cells (Fig. S7E-F), with overall enrichment that slightly exceeds that of H3K27ac (Fig. S7G). Notably, Kacme signal showed CTCF orientation-dependent asymmetry around CTCF motifs, suggesting a functional relationship between Kacme deposition and CTCF-mediated chromatin organization (*81*). This asymmetry may be reflective of CTCF-related mechanisms such as insulation (*82*), enhancer-promoter looping (*83, 84*), or the recruitment of cofactors like cohesin (*85*). Although CTCF-binding sites are largely conserved across cell types, CTCF occupancy can be cell type-specific (*86*). We performed CTCF CUT&RUN in HEK293T cells and found that Kacme levels are predictive of CTCF enrichment (Fig. 5E-F). Specifically, CTCF motifs with the highest levels of Kacme exhibited the strongest CTCF CUT&RUN signal, whereas motifs lacking Kacme enrichment showed little to no CTCF binding. These results are consistent with a model in which Kacme opposes heterochromatin spreading, potentially by promoting open chromatin and facilitating CTCF binding.

The dynamic regulation of chromatin organization and long-range looping is well characterized in the THP-1 differentiation system (*47, 49, 55*). As observed at the *CLIP2* locus, differentiation-induced loss of Kacme correlated with decreased CTCF binding and a corresponding spread of H3K27me3 and H3K9me3 (Fig. 5G). Integration with matched Hi-C data (*49*) revealed that loci that gained chromatin loops during THP-1 differentiation exhibited significantly increased promoter Kacme levels (Fig. 5H). Changes in Hi-C looping (+/- PMA) were strongly correlated with changes in Kacme levels (Fig. S7H), supporting a broader role for Kacme in genome organization.

### Kacme-enriched boundary elements preferentially resist H3K27me3 spreading following KDM6A/B inhibition

Certain chromatin-associated factors, including histone acetylation marks (e.g., H4Kac) and bromodomain-containing proteins such as BRD2, are thought to support boundary function at heterochromatin-euchromatin transition zones (*87, 88*). The enrichment of Kacme at the borders of broad H3K27me3 domains suggests that Kacme acts in concert with CTCF to oppose heterochromatin spreading. To investigate whether Kacme enrichment at CTCF boundary sites contributes to insulator activity, we first used CTCF-binding sites as genomic anchors to identify junctions with sharp transitions between active and inactive chromatin domains (*80, 89*). We identified a subset of CTCF peaks (n = 3,529) in HEK293T cells that demarcate transitions in H3K27me3 levels (Fig. 6A) and applied k-means clustering to Kacme ChIP-seq signal across these regions, revealing four distinct groups (Fig. 6B). Clusters 1 and 2 (n = 122 and n = 524 boundaries, respectively), which displayed the highest levels of Kacme enrichment, also showed the strongest co-enrichment for BRD2 (Fig. 6C) and the most pronounced restriction of H3K27me3 into flanking euchromatin (Fig. S8A), supporting a model in which Kacme strengthens CTCF-associated boundary elements to insulate active chromatin from encroaching heterochromatin.

**Fig. 6.**
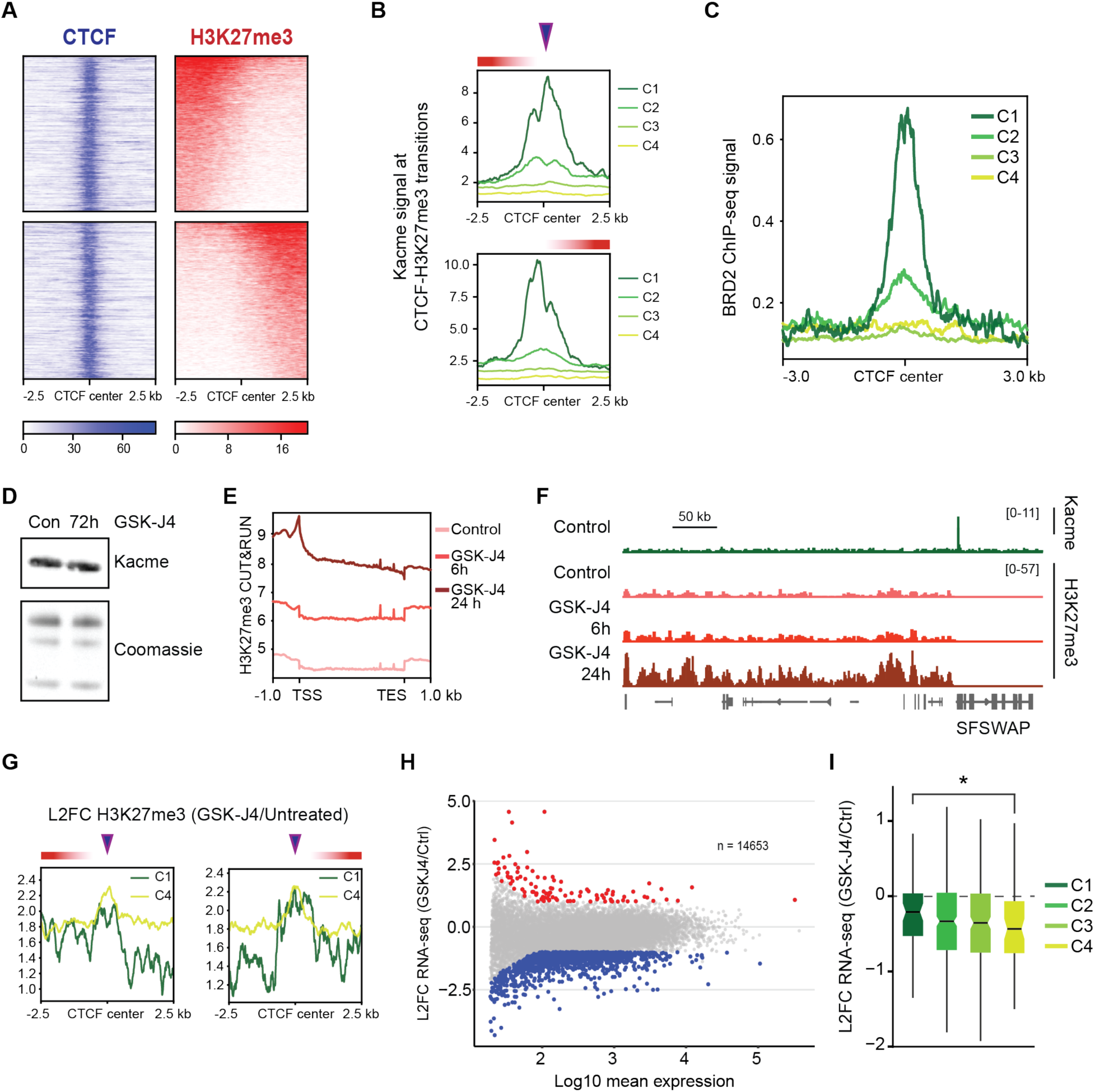
Kacme-marked boundary elements restrict H3K27me3 spreading and protect local transcription upon KDM6A/B inhibition. (**A**) Heatmaps showing CTCF and H3K27me3 signal intensity at CTCF binding sites that demarcate transitions in H3K27me3-enriched heterochromatin domains (L2FC H3K27me3 > 1). (**B**) Kacme ChIP-seq enrichment centered at CTCF-H3K27me3 transition sites (Fig. 6A), clustered by Kacme levels (C1-C4). Upper and lower plots show Kacme enrichment profiles associated with transitions out of (upper) or into (lower) H3K27me3 domains. Purple arrowheads denote the center of the CTCF-anchored transition. (**C**) BRD2 ChIP-seq signal centered on CTCF sites for the four CTCF-H3K27me3 transition clusters defined in Fig. 6B. Transitions enriched for Kacme (C1) show the strongest co-enrichment for BRD2. (**D**) Representative western blot of global Kacme levels following 72 h of GSK-J4 treatment (2.5 μM). (**E**) Metaplots of spike-in normalized H3K27me3 CUT&RUN signal in DMSO-treated HEK293T cells and HEK293T cells treated with 5 μM GSK-J4 for 6 h or 24 h. Inhibition of KDM6A/B results in a global increase in gene body H3K27me3 levels. (**F**) Genome browser tracks showing Kacme profiles in control HEK293T cells and H3K27me3 profiles in control, GSK-J4 (6 h), or GSK-J4 (24 h)-treated HEK293T cells. GSK-J4 treatment leads to increased H3K27me3 levels; however, H3K27me3 spreading does not extend beyond the Kacme-marked boundary into the *SFSWAP* locus. (**G**) L2FC in spike-in normalized H3K27me3 levels (24 h GSK-J4 vs. control) centered on CTCF transition sites (C1 and C4, as defined in Fig. 6B). Purple arrowheads denote the center of the CTCF-anchored transition. (**H**) MA plot displaying L2FC in spike-in normalized RNA expression levels (24 h GSK-J4 vs. control) versus mean expression. Red points indicate significantly upregulated genes (L2FC > 1, FDR-adjusted p < 0.05) and significantly downregulated genes (L2FC < −1, FDR-adjusted p < 0.05). (**I**) Box plots showing the L2FC in RNA expression (24 h GSK-J4 vs. control) at genes proximal to CTCF transition sites (within 10 kb of C1-C4 transition sites, as defined in Fig. 6B). Statistical significance was determined using a one-way ANOVA followed by Tukey’s HSD post-hoc test. * = p < 0.05.

To test this hypothesis, we asked whether Kacme-marked loci maintain heterochromatin-euchromatin boundary integrity under pharmacological conditions that promote heterochromatinization. We treated HEK293T cells with GSK-J4, a selective dual inhibitor of the H3K27 demethylases KDM6A/B (*90*). GSK-J4 treatment resulted in a dose- and time-dependent increase in H3K27me3 levels (Fig. S8B-C). Global Kacme levels remained largely unchanged even after 72 hours of KDM6A/B inhibition (Fig. 6D). We then performed spike-in normalized CUT&RUN against H3K27me3 in HEK293T cells treated with GSK-J4 for 0, 6, or 24 hours. Differential binding analysis revealed an increase in H3K27me3 levels across both promoters and gene bodies (Fig. 6E and Fig. S8D), validating successful KDM6A/B inhibition. While GSK-J4 treatment induced the accumulation of H3K27me3 across broad heterochromatin domains, H3K27me3 spreading was constrained at heterochromatin-euchromatin boundaries highly marked by Kacme (Fig. 6F). Analysis across the previously defined CTCF transition classes confirmed that junctions with strong Kacme enrichment exhibited significantly attenuated H3K27me3 spreading compared to Kacme-depleted boundaries (Fig. 6G).

We next determined whether Kacme-marked regions that resist H3K27me3 spreading also protect against transcriptional downregulation. To address this, we performed TimeLapse-seq in HEK293T cells treated with GSK-J4 for 24 hours, which led to clear treatment-induced changes in transcription (Fig. S8E) and the expected widespread transcriptional downregulation (Fig. 6H). Integration with the previously defined Kacme-CTCF transition clusters (Fig. 6B) revealed that genes located near Kacme-enriched insulator sites exhibited the least transcriptional downregulation and were significantly more protected than those near insulator sites lacking Kacme (Fig. 6I). Together, these results demonstrate that Kacme-enriched boundaries resist heterochromatin spreading, further establishing Kacme as a modification that maintains open chromatin at active regulatory regions.

## Discussion

Kacme is a recently discovered histone modification initially linked to increased rates of transcriptional initiation. Here, we extend these findings, revealing that Kacme is present at regions of accessible chromatin, including active promoters, enhancers, silencers, and boundary elements. Our results support a model in which Kacme functions to help sustain local chromatin accessibility, even under conditions of widespread chromatin condensation (Fig. 7).

**Fig. 7.**
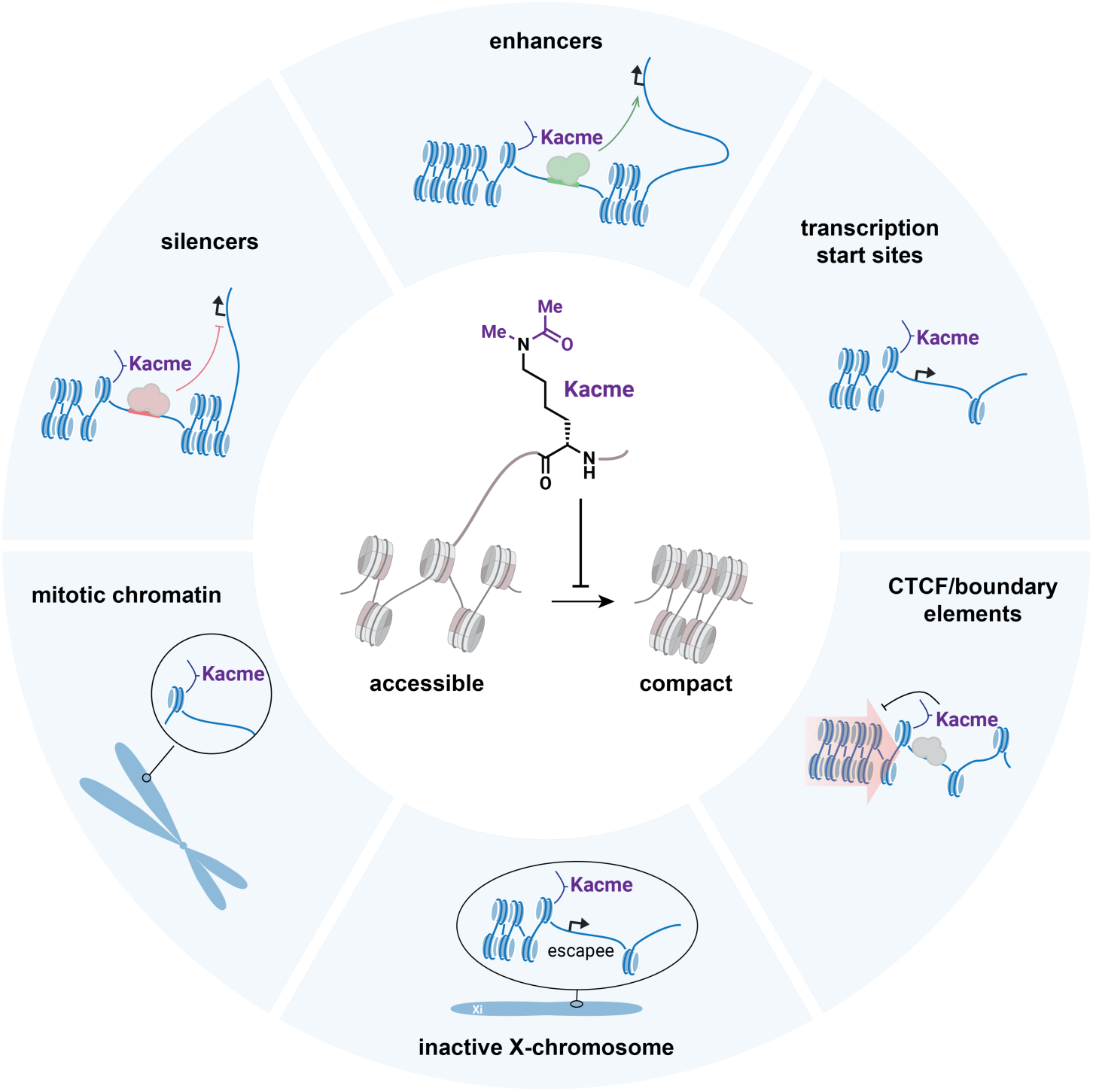
Model illustrating the proposed role of Kacme across diverse genomic contexts. Kacme is enriched at active regulatory elements, including transcription start sites, enhancers, silencers, and CTCF-bound boundary elements, where it supports local chromatin accessibility. Kacme is also retained at select loci during mitosis and on the inactive X chromosome, marking regions that resist compaction. Collectively, these observation support a unifying model in which Kacme acts to oppose chromatin compaction and preserve accessible chromatin states.

In fly and human cells, Kacme marks the TSSs of actively transcribed genes (*26*) as well as additional classes of regulatory elements (*27*). Consistent with these roles, we find that Kacme is generally associated with chromatin accessibility, independent of canonical activating marks such as H4K5ac and H3K27ac. We identified a subset of highly accessible enhancer regions in HEK293T cells that showed preferential enrichment of Kacme but not H3K27ac and H3K4me1. These enhancers were significantly overrepresented for pioneer TF motifs, including members of the FOX family, suggesting that Kacme may play a role in enhancer priming and the early establishment of chromatin accessibility (*91, 92*). Additional support for this model comes from recent evidence that during transdifferentiation of IMR90 fibroblasts into skeletal muscle progenitor cells, changes in chromatin accessibility at MYOD-bound super-enhancers most closely correlate with changes in Kacme levels (*27*). Kacme may act upstream of H3K27ac and H3K4me1 at select enhancers, either to initiate enhancer activation or reinforce chromatin accessibility following pioneer TF engagement. Future studies will be required to delineate the functional differences between Kacme-enriched and canonical active enhancers and to assess whether Kacme contributes to higher-order chromatin organization, including enhancer-promoter looping (*93*).

Several lines of evidence support a model in which Kacme acts to maintain regions of open chromatin and resist chromatin compaction. Sites preferentially enriched for Kacme exhibit hallmarks of accessible chromatin, including high ATAC-seq signal, CpG hypomethylation, anti-correlation with H3K9me3 and H3K27me3, and enrichment for TF motifs associated with pioneer activity. The accessibility of Kacme-marked chromatin is not due to ongoing transcription, as global Kacme levels remain largely unchanged following acute transcriptional inhibition. This contrasts with reports of other activating histone marks that decrease rapidly upon transcriptional inhibition (*36*). Together with previous findings linking Kacme to transcriptional initiation but not promoter-proximal pause-release (*26*), our results point to a role for Kacme in maintaining accessibility at TSSs upstream of Pol II recruitment, perhaps through mechanisms similar to those operating at enhancers and boundary elements.

To test the model that Kacme marks chromatin that resists compaction, we examined its genomic distribution in three systems characterized by widespread chromatin condensation. We found that Kacme is maintained at a subset of loci during mitosis, marking regions that retain accessibility and are associated with rapid transcriptional reactivation in early G1. In this respect, Kacme behaves similarly to H4K5ac and H3K27ac, established marks of mitotic bookmarking.

Consistent with this model, Kacme also marks genes, including *XIST*, that are expressed from or escape inactivation on the largely heterochromatic inactive female X chromosome. Further supporting this idea, Kacme is enriched at heterochromatin boundaries. Its presence at barrier regions, together with co-enrichment of CTCF and BRD2, suggests that Kacme contributes to the anti-repressive activity of insulator elements. This conclusion was reinforced by experiments in which we pharmacologically elevated H3K27me3 levels and compared CTCF boundary sites with or without Kacme. Regions marked by Kacme more effectively resisted heterochromatin spreading and better preserved expression of nearby genes.

Taken together, these diverse systems support a unified model of Kacme function: it marks regions to maintain accessibility and prevent compaction. This model accounts for Kacme’s presence at TSSs, where it contributes to promoter accessibility; at enhancers, where it maintains DNA accessibility downstream of pioneer factor activity; at loci that must remain accessible during mitosis; at *XIST* and escapee genes on the Xi; and at CTCF binding sites and heterochromatin boundaries. Mechanistically, Kacme may act through multiple, non-mutually exclusive mechanisms, including (1) promoting accessibility through lysine charge neutralization, (2) recruiting or activating complexes with Kacme readers such as BRD2, and (3) resisting HDAC-mediated removal by specific HDAC-containing complexes. Future work will be needed to dissect the relative contributions of these and other mechanisms to Kacme’s role in sustaining chromatin accessibility and opposing compaction.

## Acknowledgments

We thank B. Lesch, P.L. Puri, and all members of the Simon Lab for their comments and valuable feedback. We thank I. Vock for improvements to the computational analysis of the sequencing data. We also thank the Yale Center for Genome Analysis for library QC and sequencing.

## Funding

National Institutes of Health grant R01GM137117 (MDS)

F31CA275096-01A1 (EB)

DeLuca Center for Innovation in Hematology Research (CLB, MDS)

## Author contributions

Conceptualization: APU, CLB, LJC, MDS

Methodology: APU, CLB, LJC, MDS

Investigation: APU, JTY, CLB, LJC, YK, EB

Visualization: APU

Funding acquisition: APU, MDS

Project administration: APU, MDS

Supervision: APU, MDS

Writing – original draft: APU, MDS

Writing – review & editing: APU, JTY, CLB, LJC, YK, EB, LK, MDS

## Data and materials availability

Requests for materials generated in this study should be directed to the lead contact. The authors will share reagents upon request. Raw data of TimeLapse-seq, ATAC-seq, ChIP-seq, and CUT&RUN libraries generated in this study are publicly available and accessible on Gene Expression Omnibus (GEO) under GEO: GSE326873. Data used in, but not generated from, this study is summarized in Table S2.

Custom scripts written for downstream analyses are freely available at https://git.yale.edu/SimonLab/Pintado-Urbanc_2026_Kacme_paper. Any additional information required to reanalyze the data reported in this paper is available from the lead contact upon request.

## Materials and Methods

### Experimental model and subject details

HEK293T cells were cultured at 37°C in DMEM containing 10% FBS and 1% penicillin-streptomycin. THP-1 cells were grown at 37°C as single cell suspensions in RPMI-1640 containing 10% FBS, 1% penicillin-streptomycin, and 0.05 mM 2-mercaptoethanol (BME). THP-1 differentiation was carried out by treating cells with 40 ng/mL PMA (phorbol 12-myristate 13-acetate, Abcam, ab120297) for 48 hrs, followed by washing away non-adherent cells with PBS and incubating the adherent cells in fresh RPMI medium for an additional 24 hrs. *D. melanogaster* S2 cells were grown at 27°C in Schneider’s Drosophila Medium supplemented with 10% FBS and 1% penicillin-streptomycin.

### Small-molecule inhibitor treatments

For transcriptional inhibition studies, HEK293T cells were treated with 10 μM triptolide (MCE, HY-32735) or 500 nM flavopiridol (Sigma Aldrich, F3055) for 1hr. Unless otherwise indicated in the figure captions, experiments targeting KDM6A/B were performed by treating HEK293T cells with 5 μM GSK-J4 (MCE, HY-15648B) for either 6 or 24 hrs.

### Antibodies

The following antibodies were used: anti-Kacme (developed with Abcam; 1:1,000 (Western blots), 2.0 μg per rxn (ChIP-seq), 0.5 μg per rxn (CUT&RUN), and 1:1,000 (IF)); anti-H4Kacme (developed with Cocalico Biologicals; 1:1,000 (Western blots) and 2.0 μg per rxn (ChIP-seq); anti-H3K27me3 (EpiCypher, 13-0055, 1:10,000 (Western blots) and 0.5 μg per rxn (CUT&RUN)); anti-H3K4me3 (EpiCypher, 13-0060, 0.5 μg per rxn (CUT&RUN)); anti-H3K27ac (Abcam, ab4729, 2.0 μg per rxn (ChIP-seq)); anti-CTCF (Active Motif, 61311, 1.0 μg per rxn (CUT&RUN)); anti-H4Kac (Abcam, ab233193, 2.0 μg per rxn (ChIP-seq)); anti-H4K5ac (Millipore, 07-327, 1:2,000 (Western blots), 2.0 μg per rxn (ChIP-seq), and 0.5 μg per rxn (CUT&RUN); anti-Histone H3 (phospho S10) (Abcam, ab47297, 1:500 (Western blots)); anti-ACA(Antibodies Inc., 15-234-0001, 1:1,000); anti-IgG negative control (EpiCypher, 13-0042, 0.5 μg per rxn (CUT&RUN)); goat anti-rabbit IgG-HRP (Invitrogen, 31460, 1:2,000 (Western blots)); Alexa Fluor AffiniPure goat anti-rabbit IgG (H+L) (Jackson ImmunoResearch, 1:1,000).

### Preparation of histone extract

Histone extracts were collected using the EpiQuik Total Histone Extraction Kit (Epigentek) following the standard kit instructions.

### Western blotting

Protein samples were boiled in 5× SDS sample buffer (0.25 M Tris pH 6.8, 10% SDS, 50% glycerol, 0.5% DTT, 0.25% bromophenol blue) for 5 mins, then resolved by SDS–PAGE using a Novex Mini Cell gel box (Invitrogen) with NuPAGE MOPS SDS Running Buffer (Invitrogen). PVDF membranes (Immobilon-P, 0.45 µm pore size, Millipore) were pre-soaked in methanol for 3 mins and equilibrated in NuPAGE Transfer Buffer (Invitrogen) for 30 mins. Protein transfer was performed using the Novex XCell II blotting system (Invitrogen) following the manufacturer’s instructions. Membranes were blocked in 5% milk in PBS-T for 1 hr at room temperature with rotation, and then incubated with primary antibody (at the indicated dilution) in 5% milk in PBS-T overnight at 4 °C. After three 5 min washes in TBS-T, membranes were incubated with secondary antibody (at the indicated dilution) in 5% milk in PBS-T. Following three 5 min washes in TBS-T and two washes in PBS-T, blots were developed using Clarity Western ECL substrate (BioRad). Imaging was performed on an ImageQuant LAS 4000 system (GE). Band intensities were quantified using ImageJ and normalized to total histone signal as detected by Coomassie or Ponceau staining.

### ATAC-seq

ATAC-seq datasets were processed using the nf-core/atacseq pipeline (version 2.1.2) (*94*). In brief, adapter sequences were trimmed using Cutadapt (*95*) and reads were aligned to the human hg38 genome using the BWA aligner (*96*). PCR duplicates and reads mapping to mitochondrial DNA were filtered out. Peaks were called using MACS2 (*97*). Normalization scale factors for ATAC-seq were calculated using TMM normalization (*98*). ATAC peaks were annotated using ChIPseeker (*99*), and those located within ±50 bp of an annotated transcription start site were classified as promoter-associated ATAC peaks. Metaplots of ATAC-seq density were created using deepTools (*100*).

### ChIP-seq

ChIP–seq was performed as previously described (*26*), with all experiments conducted in duplicate. For each immunoprecipitation, ∼20 × 10⁶ HEK293T or THP-1 cells were used. Cells were washed twice with PBS and cross-linked with 1% formaldehyde (Pierce) for 10 mins at room temperature, followed by quenching with 125 mM glycine for 5 mins and two additional PBS washes.

Cells were lysed in 4 ml lysis buffer (50 mM HEPES pH 7.9, 140 mM NaCl, 1 mM EDTA, 10% glycerol, 0.5% NP-40, 0.25% Triton X-100, 1× protease inhibitors) for 10 mins on ice, centrifuged at 4,000 rpm for 5 mins at 4 °C, and washed twice with 4 ml cold wash buffer (10 mM Tris-HCl pH 7.5, 200 mM NaCl, 1 mM EDTA pH 8.0, 0.5 mM EGTA pH 8.0, 1× protease inhibitors). Pellets were resuspended in 1 ml shearing buffer (0.1% SDS, 1 mM EDTA, 10 mM Tris pH 7.5, 1× protease inhibitors) and sheared with a Covaris S220 (140 W, 5% duty factor, 200 cycles/burst, 4 °C) for 12 mins (HEK293T) or 24 mins (THP-1), as determined by time-course optimization.

After the addition of Triton X-100 (1% final) and NaCl (150 mM final), the extract was cleared by centrifugation at 21,000Xg for 10 mins at 4 °C, and 10% was set aside as input. Remaining extract was incubated overnight at 4 °C with 2 µg antibody. Immunoprecipitated chromatin was captured on 30 µl Dynabeads Protein G (Invitrogen) at 4 °C for 1.5 hrs. Beads were washed twice with low-salt buffer (0.1% SDS, 1% Triton X-100, 2 mM EDTA, 20 mM HEPES-KOH pH 7.5, 150 mM NaCl), twice with high-salt buffer (as above but 500 mM NaCl), once with LiCl buffer (100 mM Tris-HCl pH 7.5, 0.5 M LiCl, 1% NP-40, 1% sodium deoxycholate), and once with TE buffer (10 mM Tris-HCl pH 8.0, 0.1 mM EDTA).

Chromatin was eluted in 100 µl proteinase K buffer (20 mM HEPES pH 7.5, 1 mM EDTA, 0.5% SDS) with 40 µg proteinase K (Ambion) for 30 mins at 50 °C. Cross-linking was reversed by adding NaCl (150 mM final) and 0.25 µg DNase-free RNase (Roche) and incubating overnight at 65 °C. DNA was purified with the QIAquick PCR Purification Kit (Qiagen) for use in qPCR or library preparation.

Libraries were prepared either by the Yale Center for Genome Analysis or following a modified published protocol (*101*). Briefly, DNA ends were polished with T4 Ligase Buffer (NEB), T4 PNK (NEB), T4 DNA Polymerase (NEB), and Klenow DNA polymerase (NEB). After isolation of DNA with the MinElute PCR Purification Kit (Qiagen), A-tailing was performed with NEB Buffer 2 and Klenow 3′-5′ exo minus (NEB). Adapter ligation was performed using Blunt/TA Ligase Mastermix (NEB), and resulting libraries were PCR-amplified for 6-14 cycles with indexing primers. Libraries were purified with 1.5× SPRI bead purification (Beckman Coulter) and sequenced on an Illumina NovaSeq 6000 2X100bp.

### ChIP-seq data analysis

ChIP-seq data was processed using the nf-core/chipseq pipeline (version 2.0.0) (*94*). In brief, adapter sequences were trimmed using Trim Galore (*95*) and reads were aligned to the human hg38 reference genome using the BWA aligner. Duplicate reads and reads mapping to blacklisted regions were filtered out. Counts Per Million (CPM) normalization was applied to create scaled bigWig files. Peaks were called with MACS2 using default parameters in narrow peak mode, with inputs used as controls. Peak-to-gene annotation was performed using ChIPseeker. Metaplots and heat maps of ChIP-seq density were created with deepTools. For heat map generation, colocalization, and pathway analyses, consensus peak sets were identified using BEDTools (*102*) to retain only peaks present in both replicates. K-means clustering in deepTools was used to compare Kacme and H3K27me3 enrichment profiles in HEK293Ts at annotated TSSs in the human hg38 reference genome.

Differential binding analysis of histone PTM ChIP-seq data was performed with DiffBind using standard parameters (*103*). Known motif analysis was performed using HOMER (*104*). For Fig. 5C-D, enriched motifs were identified within the regions ±500 bp of peak centers with log_2_[Kacme/H3K27ac] > 0 and FDR < 0.01 (as determined by DiffBind from HEK293T ChIP-seq data).

### Analysis of Kacme enrichment at enhancer and silencer regions

HEK293T-specific enhancer regions were obtained from the EnhancerAtlas 2.0 database (*37*). To distinguish enhancer-associated Kacme enrichment from promoter-associated Kacme enrichment, enhancer regions within ±3 kb of H3K4me3 peaks were excluded. HEK293T H3K4me3 consensus peaks were identified as regions with overlapping MACS2-called peaks across two biological replicates. After distance-based filtering with BEDTools, 8,477 high-confidence promoter-distal enhancers were identified from the original set of 21,506 enhancer regions. K562-specific silencer elements were obtained from (*105*). K-means clustering (k=2) was performed on K562 ATAC-seq signal at these regions. Kacme ChIP-seq and ATAC-seq signal were plotted for the two resulting clusters.

### ChromHMM and HOMER motif analysis

ChromHMM (*41*) was used for chromatin-state discovery in K562 cells. Peak-calling BED files for 16 histone post-translational modifications, histone variants, and chromatin-associated factors were obtained from the ENCODE Project and in-house ChIP-seq experiments. BED files were binarized using the BinarizeBed command with hg38 as the reference genome and the - center option. The LearnModel command was executed with 15 chromatin states specified. Motif enrichment analysis was performed on genomic regions corresponding to ChromHMM state E12, using the findMotifsGenome.pl function in HOMER, with hg38 as the reference genome and a 200 bp window centered on each interval.

### RRBS data analysis

Aligned reduced representation bisulfate sequencing (RRBS) data from HEK293s, available through the ENCODE Project Consortium, was analyzed to assess CpG methylation at consensus Kacme-marked regions (± 20 kb). To control for the presence of other activating acetylation marks, CpG methylation levels were examined at the top 2000 peak regions with the highest relative Kacme-to-H3K27ac signal ratios, as quantified by DiffBind (see *ChIP-seq data analysis* section). All profile plots were made using deepTools.

### RT-qPCR analysis of differentiated THP-1 cells

Total RNA from THP-1 cells treated with either DMSO or PMA was extracted using TRIzol reagent and precipitated with isopropanol. Genomic DNA was removed with TURBO DNase, followed by RNA purification with an equal volume of Agencourt RNAClean XP beads. 1 μg of RNA was reverse-transcribed into cDNA using the SuperScript VILO cDNA Synthesis Kit (Invitrogen). Quantitative PCR (qPCR) was performed on cDNA using iTaq Universal SYBR Green Supermix (Bio-Rad). GAPDH was used as a normalization control, and relative gene expression was calculated using the ΔΔCt method. Primers for targets are included in Table S1.

### ChIP-seq, ATAC-seq, RNA-seq, and Hi-C analysis of differentiated THP-1 cells

Literature-sourced ChIP-seq and ATAC-seq data from undifferentiated and differentiated THP-1 cells were processed as described above (see *ATAC-seq* and *ChIP-seq data analysis*). DiffBind was used to assess significant differences in histone PTM enrichment levels. Normalization and differential expression analysis of RNA-seq performed in PMA-treated cells (data obtained from (*55*)) was performed using DESeq2.

Hi-C data for undifferentiated and differentiated THP-1 cells was obtained from a previously published study (*49*), in which genomic regions were classified as having “significantly gained”, “significantly lost”, or “unchanged” looping upon differentiation. Hi-C looping regions were annotated to the nearest gene using ChIPseeker. Violin plots were used to evaluate differentiation-induced changes in promoter-proximal Kacme ChIP-seq signal (±3 kb from the TSS) at genes with significant gains in looping. Statistical significance was assessed via Wilcoxon rank-sum tests.

### CUT&RUN

CUT&RUN was performed in duplicate in HEK293T nuclei using the EpiCypher CUTANA ChIC/CUT&RUN Kit following the standard protocol. For Kacme, H4K5ac, and CTCF CUT&RUN samples, cells were crosslinked (0.1% formaldehyde, 1 min) prior to nuclei isolation. For GSK-J4 CUT&RUN experiments, *E. coli* spike-in DNA was included to be able to normalize for global changes in H3K27me3 levels. Library preparation was carried out with the EpiCypher CUTANA CUT&RUN Library Prep Kit, and sequencing was performed on a NovaSeq 6000 system with 2X100bp reads. Adapter sequences were trimmed using fastp (*106*), and the reads were aligned to the human hg38 genome using bwa-mem2 (*107*). Aligned reads were processed using SAMtools (*108*), and profile plots and heatmaps were generated using deepTools. Peak calling was performed using MACS2 in broad peak mode for H3K27me3 samples and in narrow peak mode for all other targets, with IgG samples used as controls.

### Mitotic arrest and release time course

Mitotic arrest and release were performed using a protocol adapted from [(*59*)]. For each sample, 2 million HEK293T cells were seeded and allowed to adhere overnight. The following day, nocodazole (Sigma, M1404) was added to the media at a final concentration of 50 ng/mL, and cells were incubated for 16 hrs at 37 °C. Mitotic cells were harvested by manual shake-off.

For mitotically arrested samples (*T0*), cells were pelleted (600Xg, 3 min, RT), washed twice with cold PBS, and resuspended in medium containing 10% DMSO.

For mitotic release samples (T40, T80, T105, T165, T300), mitotic cells were harvested by shake-off, pelleted (600Xg, 3 min, RT), washed twice with room-temperature PBS, and resuspended in pre-warmed medium. To release the cells from arrest, washed cell suspensions were transferred to new plates with fresh medium and incubated at 37 °C until collection at the indicated time points. All samples were slow-frozen at −80 °C.

### Kacme immunofluorescence

HeLa cells were arrested in 100 ng/mL nocodazole for 4 hrs. Mitotic cells were isolated by shake off, washed in PBS, and resuspended in 75 mM KCl for 15 mins on ice. Cells were then spun onto glass slides. Slides were blocked in PBS + 1% BSA for 10 mins before incubation with primary antibodies diluted in PBS + BSA in a humidified chamber on ice for 4 hrs. Following primary incubation and washing, slides were incubated in fluorophore-conjugated secondary antibodies and DAPI diluted in PBS + 1% BSA for 1 hr. After a brief wash, slides were fixes in 4% paraformaldehyde for 15 mins before mounting with Prolong Gold (Invitrogen). All images were acquired on a Nikon ECLIPSE Ti2 widefield epi-fluorescence microscope with a Hamamatsu Fusion sCMOS camera. Image acquisition was managed using NIS-Elements (Nikon). All images were analyzed in FIJI (ImageJ).

### Mitotic release CUT&RUN analysis

TMM normalization was performed to obtain normalization factors, which were used for generating scaled bigWig files and for downstream differential binding analyses. Metaplots and heat maps of CUT&RUN signal at TSSs, HEK293T-specific enhancer regions, and CTCF-binding sites were created with deepTools. Peak-to-gene annotation was performed using ChIPseeker. Differential binding analysis of Kacme and H4K5ac CUT&RUN signal across timepoints was carried out using DiffBind. Genes with significantly differential Kacme at promoters (±3 kb of annotated TSSs) in mitotic (T0) and asynchronous cells were defined as those with |L2FC| > 1 and p-value ≤ 0.05.

To determine the subset of genes most strongly marked by Kacme and/or H4K5ac during mitosis, TSS-proximal peak enrichment (±1 kb of annotated TSSs) was quantified across all mitotic release time points (T0, T40, T80, T105, T165, T300) using DiffBind outputs. For genes with multiple TSS-proximal peaks, the peak with the highest signal was retained. Normalized CUT&RUN signal at these peak regions was then assembled into time-course matrices. Hierarchical clustering was performed separately on the Kacme and H4K5ac time-course matrices using the default settings of the R pheatmap package (*109*), yielding four clusters (C1-C4) for each mark that reflect distinct patterns of mitotic enrichment. These gene clusters were used for all subsequent integrative analyses with matched SF-TL-seq and ATAC-seq datasets. Transcription factor enrichment analysis on the set of genes with the highest mitotic-Kacme signal (C4, n = 1,579) was performed in Enrichr using the ENCODE TF ChIP-seq 2015 dataset. For Fig. 3J, genes in Kacme C4 but not H4K5ac C4 (highest mitotic H4K5ac signal; n = 1,645) were classified as Kacme-specific marked, those in H4Kac C4 but not Kacme C4 as H4K5ac-specific marked, and genes present in both Kacme C4 and H4K5ac C4 clusters as co-marked.

Differential binding analysis of Kacme and H4K5ac CUT&RUN data specifically at mitosis (T0) was performed to identify genes with preferential Kacme enrichment at promoters (±3 kb of annotated TSSs). Genes with significantly higher relative Kacme signal were defined by |L2FC| > 0 and p-value ≤ 0.05. The resulting set of 874 genes were analyzed in Enrichr with the Reactome Pathways 2024 dataset. For Fig. S6F, genes were classified as in Fig. 3F.

### Mitotic release TimeLapse-seq

Short-feed TimeLapse-seq experiments were done in triplicate in HEK293T cells released from mitosis. Metabolic labeling was performed essentially as previously described (*66*). Cells were spiked with s^4^U (500 μM) (Thermo Fisher) for the last 20 mins prior to sample collection. After the labeling period, plates were immediately placed on ice and washed with ice-cold PBS. Cells were scraped, pelleted by centrifugation at 500Xg for 5 mins, resuspended in 1 mL of TRIzol, and frozen at −80°C. Total RNA was isolated and extracted with phenol-chloroform and precipitated in isopropanol supplemented with 1 mM DTT. After genomic DNA depletion with TURBO DNase, total RNA was isolated with one equivalent volume of Agencourt RNAClean XP beads. 5 μg of total RNA was subjected to TimeLapse chemistry using meta-chloroperoxybenzoic acid (Alfa Aesar) as the oxidant. 10 ng of TimeLapse-treated RNA input was used to prepare sequencing libraries using the SMARTer Stranded Total RNA-Seq Kit v2 (Pico Input) with ribosomal cDNA depletion. Paired-end sequencing was performed on a NovaSeq 6000 2X150bp. Sequencing data was processed using the fastq2EZbakR pipeline (*110*). In brief, adapter sequences were trimmed using fastp and reads were aligned to the human hg38 genome using STAR. T-to-C mutations corresponding to newly transcribed RNA reads were called with a custom Python script, and the final mutation counts and feature assignments were summarized in a cB output file.

New RNA reads inferred from the TimeLapse-seq data at mitotic-release timepoints (T0, T40, T105, T300, and asynchronous) were analyzed to investigate nascent transcriptional reactivation upon mitotic exit. Normalization scale factors were calculated based on the total library size of each sample. New read counts were first normalized to reads per million (RPM) by dividing by the respective library size in millions. To account for potential gene length biases, RPM values were further normalized to reads per kilobase per million mapped reads (RPKM), providing length-adjusted expression estimates.

For each gene, RPKM values at each timepoint were normalized to the corresponding asynchronous RPKM level, yielding expression as a fraction of the ultimate asynchronous expression. Following previous approaches (*59*), genes were manually clustered based on the earliest timepoint at which their expression reached at least 50% of asynchronous levels, defining four reactivation groups: early (T40), intermediate (T105), late (T300), and asynchronous (Asyn). Z-score scaling was applied across genes to center and standardize expression values. Genes were then reordered by cluster membership and within-cluster z-score ranking to visually depict waves of transcriptional reactivation after mitosis. Heatmaps were generated using the *pheatmap* package.

RPKM-normalized expression values were plotted for each mitotic-marking cluster (see *Mitotic release CUT&RUN analysis*), enabling correlation of transcriptional reactivation kinetics with chromatin enrichment patterns. For Fig. S6D, in order to visualize reactivation dynamics relative to early and fully reactivated states, median RPKM values for each bookmarking cluster were further scales using the formula: 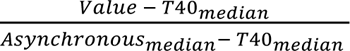 This normalization adjusts expression levels at the earliest post-mitotic timepoint (T40) to zero and asynchronous levels to one, allowing direct comparison of reactivation kinetics across clusters.

### Allele-specific ChIP-seq analysis

Curated single-nucleotide polymorphism (SNP) data for HEK293T cells were obtained from Complete Genomics whole-genome sequencing datasets (release v2; http://bioit2.irc.ugent.be/prx/hek293/v2/index.php) (*75*). To enable allele-specific analyses, variants were filtered using bcftools (*111*) to retain only heterozygous, biallelic SNPs. Bulk Kacme ChIP-seq BAM files were then filtered for reads overlapping the curated SNPs on the X chromosome and partitioned into allele-specific read sets using a custom Python script implemented with the *pysam* library. For each heterozygous SNP, overlapping reads were identified, with unmapped and duplicate reads excluded. The aligned base at the SNP position was extracted from each read and compared to the reference and alternate alleles; reads containing the reference or alternate allele were written to corresponding allele-specific BAM files, while reads with non-matching bases or indels at the variant position were excluded. For *XIST* and *DDX3X,* this strategy enabled direct assignment of individual ChIP-seq reads to their respective alleles based on nucleotide identity at curated heterozygous SNP positions; the same approach was applied to RNA-seq data from untreated HEK293T cells. Allele-specific read counts were computed using the GATK ASEReadCounter tool (*112*).

### Broad H3K27me3 domain identification

Broad H3K27me3 heterochromatin domains were defined as follows. H3K27me3 peaks were called from CUT&RUN performed in two biological replicates of HEK293T cells using MACS2 with the --broad peak calling option. For each replicate, H3K27me3 peak regions were iteratively merged using BEDTools (merge -d 75000) until no intervals within 75 kb could be further joined. Consensus H3K27me3 domains were obtained by intersecting the merged replicate files with BEDTools (intersect -wa). Additional filtering was performed in R to retain only domains greater than 8 kb in length. The final filtered bed file consisted of 4,162 broad H3K27me3 domains.

### CTCF and Kacme co-localization analyses

CTCF binding sites predicted for the human genome (hg38) were obtained from the FIMO-predicted dataset (AnnotationHub, JASPAR 2022, “AH104729”) in the *CTCF* Bioconductor package (*113*). Predicted binding sites were filtered to retain only those with a motif *p*-value < 1×10^-6^ and restricted to standard chromosomes. The filtered dataset contained 21,671 sites, which were then sorted and strand-annotated to distinguish forward (“+”) and reverse (“–”) orientations. K-means clustering (k=4) was performed on HEK293T Kacme ChIP-seq signal at these regions, and HEK293T CTCF CUT&RUN signal was subsequently plotted for each resulting cluster. Metaplots and clustering were performed with deepTools.

Consensus CTCF-binding sites in HEK293T cells were identified from two CUT&RUN biological replicates. CTCF-heterochromatin transition sites were defined as those that marked a transition in total H3K27me3 enrichment between the regions 2.5 kb upstream and 2.5 kb downstream of the CTCF binding site, with a |L2FC| > 1, adapted from [(*89*)]. K-means clustering (k=4) was performed on HEK293T Kacme ChIP-seq signal at these transition regions. H3K27me3 CUT&RUN signal was plotted for each resulting cluster.

### GSK-J4 CUT&RUN and RNA-seq

H3K27me3 CUT&RUN was performed in HEK293T cells treated with DMSO or 5 μM GSK-J4 for 6 or 24 hrs. To account for global changes in H3K27me3 levels, spike-in normalization was conducted using *E. coli* DNA read counts. Normalization factors were calculated as the reciprocal of the proportion of *E. coli* spike-in reads per sample. Spike-in normalized signal tracks and metaplots over gene bodies and TSSs were generated using deepTools. Log2 Fold Change enrichment tracks (24hr GSK-J4 vs. Untreated) were generated using the deepTools bigwigCompare function, and visualized over predefined CTCF-heterochromatin clusters (see *CTCF and Kacme co-localization analyses*, Fig. 6B).

Matched TimeLapse-seq was performed in cells treated with DMSO or 5 μM GSK-J4 for 24 hrs and spiked with s^4^U (500 μM) (Thermo Fisher) for the last 20 mins prior to sample collection. Sample collection followed the same protocol as in [(*66*)], and bioinformatic processing was performed as described above (see *Mitotic release TimeLapse-seq*). DESeq2 was used to calculate fold changes in total RNA reads after GSK-J4 treatment, filtering for genes with at least 20 reads across all samples. Normalization scale factors were calculated with DESeq2 using read counts from *Drosophila* spike-in RNA (using the accessor function sizeFactors()). CTCF-heterochromatin transition sites (see *CTCF and Kacme co-localization analyses*, Fig. 6B) were filtered for those located within 10 kb of an annotated TSS, annotated to their nearest gene, and integrated with gene expression fold changes following 24 hr GSK-J4 treatment.

---

**Fig. S1.**
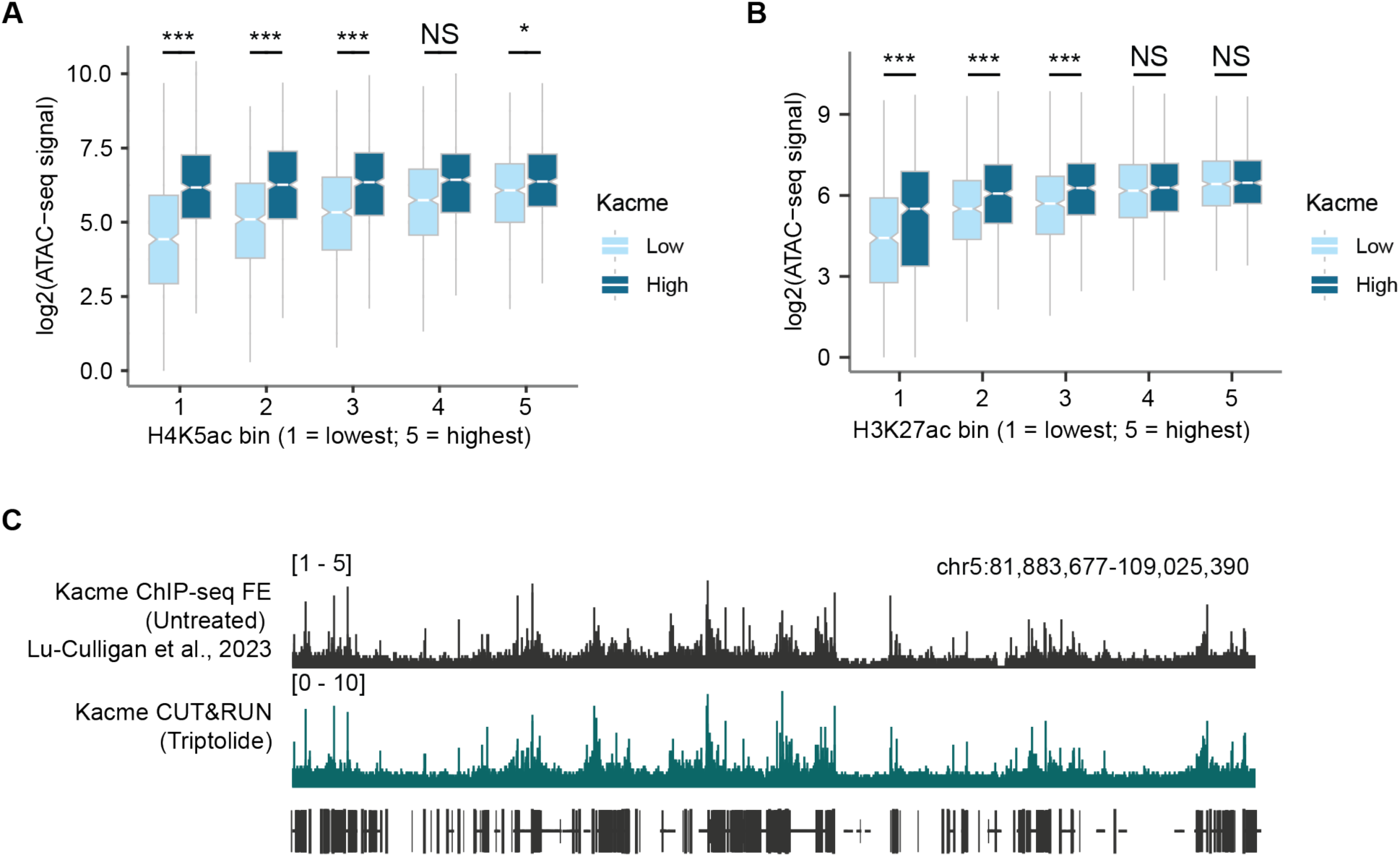
(**A**) Relative promoter accessibility in THP-1 cells for genes in the 25^th^ or 75^th^ percentiles of Kacme ChIP-seq levels. Data are binned by H4K5ac ChIP-seq signal. (**B**) Relative promoter accessibility in THP-1 cells for genes in the 25^th^ or 75^th^ percentiles of Kacme ChIP-seq levels. Data are binned by H3K27ac ChIP-seq signal. (**C**) Genome browser visualization of Kacme ChIP-seq (untreated) and Kacme CUT&RUN (triptolide-treated) profiles at representative loci on chromosome 5 in HEK293T cells. (**B-C**) Distribution means compared with two-tailed unpaired Wilcoxon test for two biological replicates. *** = p < 0.001; * = p < 0.05; NS = Not Significant.

**Fig. S2.**
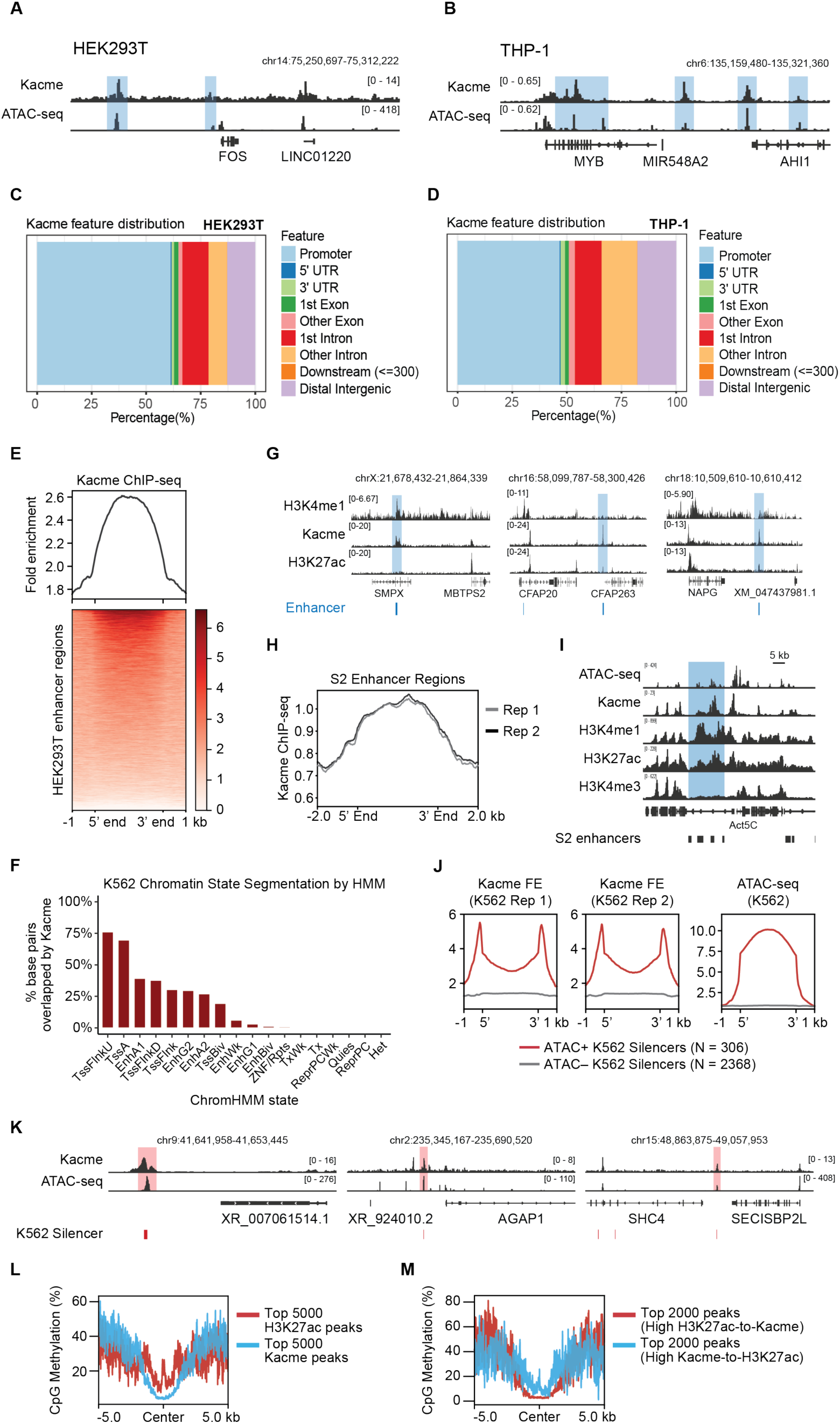
**(A-B)** Genome browser tracks showing Kacme ChIP-seq and ATAC-seq signal profiles at representative TSS-distal regions (highlighted in blue) in (A) HEK293T and (B) THP-1 cells. **(C-D)** Genomic feature distribution of consensus Kacme ChIP-seq peaks in (C) HEK293T and (D) THP-1 cells. Peaks were annotated using the ChIPseeker Bioconductor package. **(E)** Heatmap showing HEK293T Kacme ChIP-seq fold enrichment across HEK293T-specific enhancer regions (± 1kb from enhancer boundaries). **(F)** Overlap analysis between Kacme peaks (called with MACS2) and ChromHMM chromatin state annotations from the Kellis laboratory 18-state model in K562 cells (ENCSR961HFL, (*39*)). Bars indicate the percentage of base pairs within each ChromHMM state that overlap Kacme peak regions. **(G)** Example genome browser tracks showing H3K4me1, Kacme, and H3K27ac ChIP-seq enrichment at annotated enhancer regions in HEK293T cells (highlighted in blue). **(H)** Profile plot showing S2 Kacme ChIP-seq enrichment across annotated S2 enhancer regions (± 2kb from enhancer boundaries). **(I)** Example genome browser track showing ATAC-seq, H3K4me1, Kacme, and H3K27ac co-occupancy at an *Act5c*-proximal enhancer region in *Drosophila* S2 cells (highlighted in blue). Notably, H3K4me3 signal is absent, consistent with the enhancer identity of this region rather than a promoter-associated state. **(J)** Profile plots of Kacme ChIP-seq enrichment and ATAC-seq signal across annotated K562 silencer elements (± 1kb), obtained from (*105*). Silencer elements were clustered by ATAC-seq signal. **(K)** Example genome browser tracks showing Kacme and ATAC-seq signal profiles at annotated silencer elements in K562 cells (highlighted in red). **(L)** Metaplot analysis of average CpG methylation levels centered on the top 5,000 Kacme or H3K27ac ChIP-seq peak summits (ranked by their respective enrichment scores) in HEK293T cells. **(M)** Metaplot analysis of average CpG methylation levels centered on the top 2,000 peaks ranked by relative H3K27ac-to-Kacme signal or Kacme-to-H3K27ac signal in HEK293T cells.

**Fig. S3.**
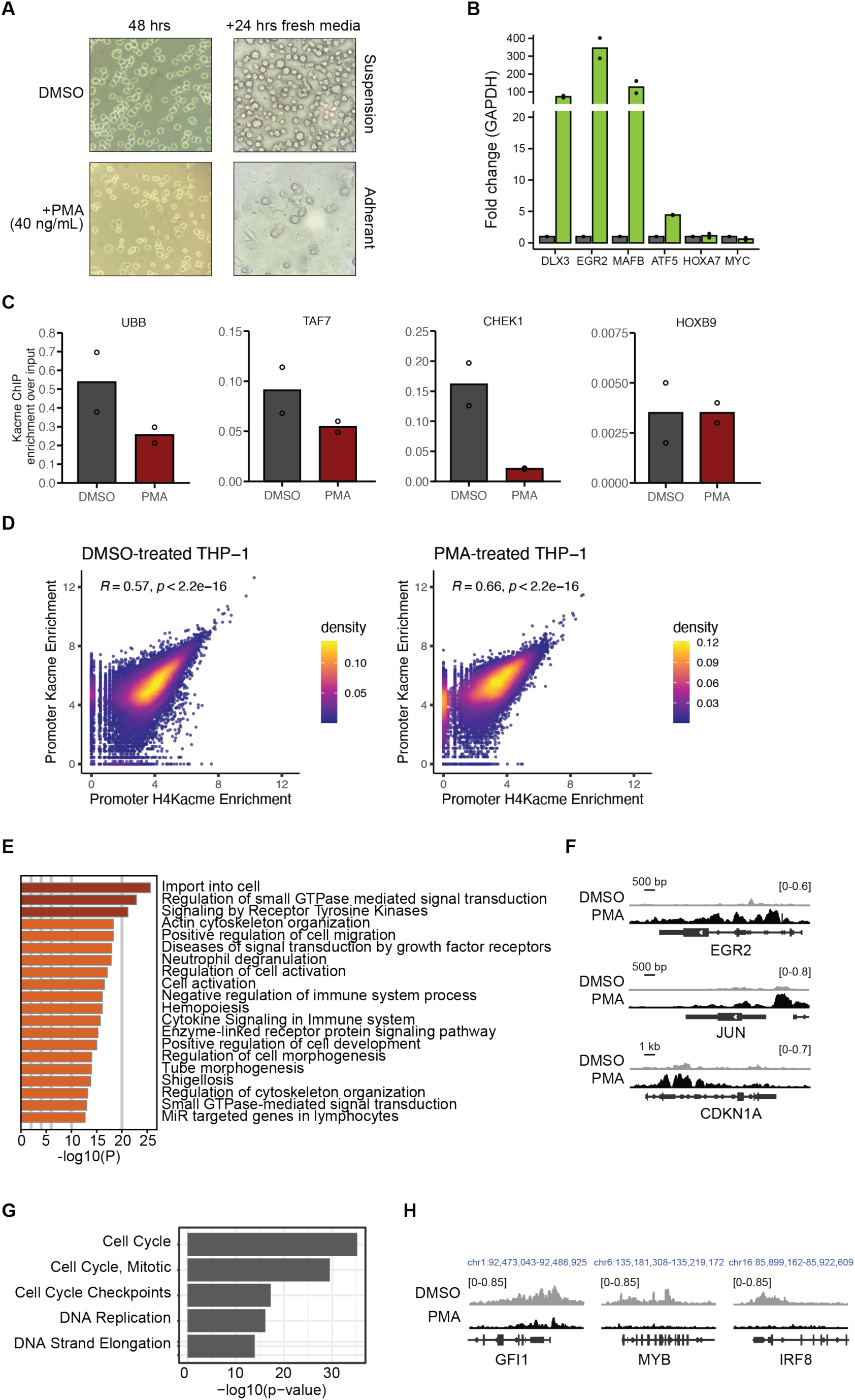
**(A)** Representative bright-field microscopy images of THP-1 cells treated with DMSO (control) or PMA (40 ng/mL). After 48 h, PMA-treated cells transition from suspension to an adherent, macrophage-like morphology. **(B)** RT-qPCR analysis showing fold change (normalized to GAPDH) of selected genes in DMSO-versus PMA-treated THP-1 cells. PMA treatment strongly induces the expression of differentiation-associated factors such as DLX3, EGR2, and MAFB. **(C)** Kacme ChIP-qPCR quantification at selected promoter regions (UBB, TAF7, CHEK1, HOXB9) in DMSO-versus PMA-treated THP-1 cells. **(D)** Genome-wide correlation of promoter H4Kacme (x-axis) and promoter Kacme (y-axis) ChIP-seq enrichment in DMSO-treated (left) and PMA-treated (right) THP-1 cells. Scatter plots show significant correlation between H4Kacme and Kacme levels, as assayed by Pearson correlation analysis. **(E)** Gene Ontology (GO) enrichment analysis for genes with significantly upregulated promoter Kacme signal (n = 2,400 genes; L2FC > 0, FDR-adjusted p < 0.05) upon PMA-induced differentiation of THP-1 cells. Top-enriched terms relate to macrophage-related processes such as cell import and neutrophil degranulation. **(F)** Genome browser tracks illustrating Kacme ChIP-seq signal at the *EGR2*, *JUN*, and *CDKN1A* loci in DMSO- and PMA-treated THP-1 cells. **(G)** Reactome Pathways 2024 enrichment analysis for genes with significantly downregulated promoter Kacme signal (n = 2,826 genes; L2FC < 0, FDR-adjusted p < 0.05) upon PMA-induced differentiation of THP-1 cells. **(H)** Genome browser tracks illustrating Kacme ChIP-seq signal at the *GFI1*, *MYB*, and *IRF8* loci in DMSO- and PMA-treated THP-1 cells.

**Fig. S4.**
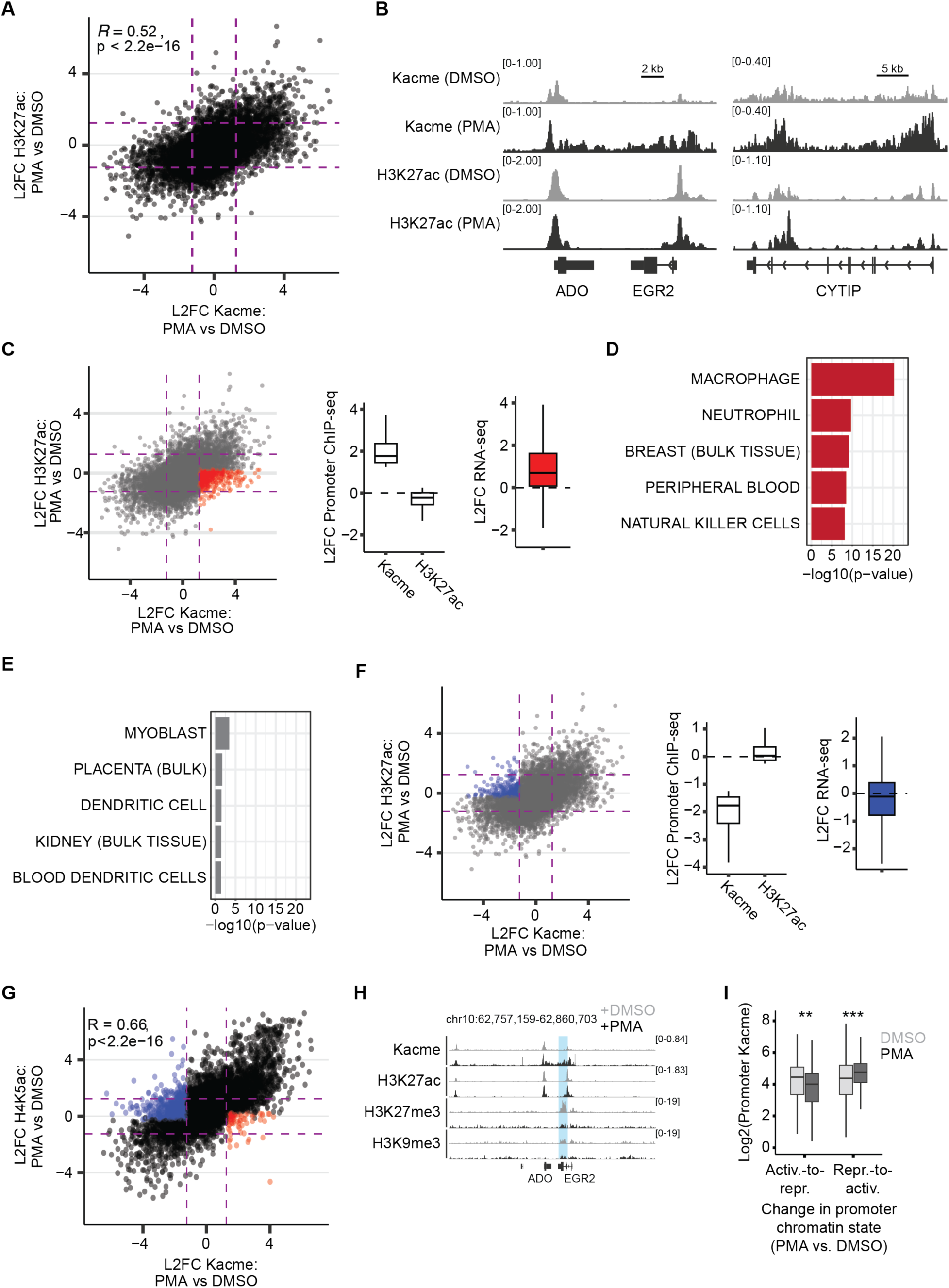
**(A)** Genome-wide comparisons of promoter-associated Kacme (x-axis) and H3K27ac (y-axis) changes upon PMA treatment in THP-1 cells. Each dot represents the mean promoter signal change for a single gene. Dashed lines represent a ± 1.25 L2FC. Pearson correlation and significance are shown. **(B)** Genome browser tracks illustrating Kacme and H3K27ac ChIP-seq signal at the *EGR2*, *ADO*, and *CYTIP* loci in DMSO- and PMA-treated THP-1 cells. **(C)** Left: Same as Fig. S4A. Genes with an average L2FC in promoter Kacme signal > 1.25 and an average L2FC promoter H3K27ac signal < 0.25 are highlighted in red. Middle: Boxplot representation of the average L2FC in promoter Kacme and H3K27ac signal for genes highlighted in red. Right: RNA-seq L2FC (PMA/DMSO) for corresponding genes. **(D)** ARCHS4 Tissues enrichment analysis for genes with an average L2FC in promoter Kacme signal > 1.25 and an average L2FC promoter H3K27ac signal < 0.25 (PMA/DMSO). **(E)** ARCHS4 Tissues enrichment analysis for genes with an average L2FC in promoter H3K27ac signal > 1.25 and an average L2FC promoter Kacme signal < 0.25 (PMA/DMSO). **(F)** Left: Same as Fig. S4A. Genes with an average L2FC in promoter Kacme signal < −1.25 and an average L2FC promoter H3K27ac signal > −0.25 are highlighted in blue. Middle: Boxplot representation of the average L2FC in promoter Kacme and H3K27ac signal for genes highlighted in blue. Right: RNA-seq L2FC (PMA/DMSO) for corresponding genes. **(G)** Genome-wide comparisons of promoter-associated Kacme (x-axis) and H4K5ac (y-axis) changes upon PMA treatment in THP-1 cells. Each dot represents the mean promoter signal change for a single gene. Dashed lines represent a ± 1.25 L2FC. Pearson correlation and significance are shown. Genes with an average L2FC in promoter Kacme signal > 1.25 and an average L2FC promoter H4K5ac signal < 0.25 are highlighted in red. Genes with an average L2FC in promoter Kacme signal < −1.25 and an average L2FC promoter H4K5ac signal > −0.25 are highlighted in blue. **(H)** Genome browser tracks illustrating Kacme, H3K27ac, H3K27me3, and H3K9me3 ChIP-seq signal at the *EGR2* and *ADO* loci in DMSO- and PMA-treated THP-1 cells. The PMA-induced increase in Kacme at this region is accompanied by a loss of heterochromatic H3K27me3. **(I)** Boxplots showing the L2FC in Kacme ChIP-seq enrichment at promoters switching between active and repressive chromatin states upon PMA treatment, as reported in (*55*). Statistical significance was determined using pairwise Wilcoxon rank-sum tests. *** = p < 0.001; ** = p < 0.01.

**Fig. S5.**
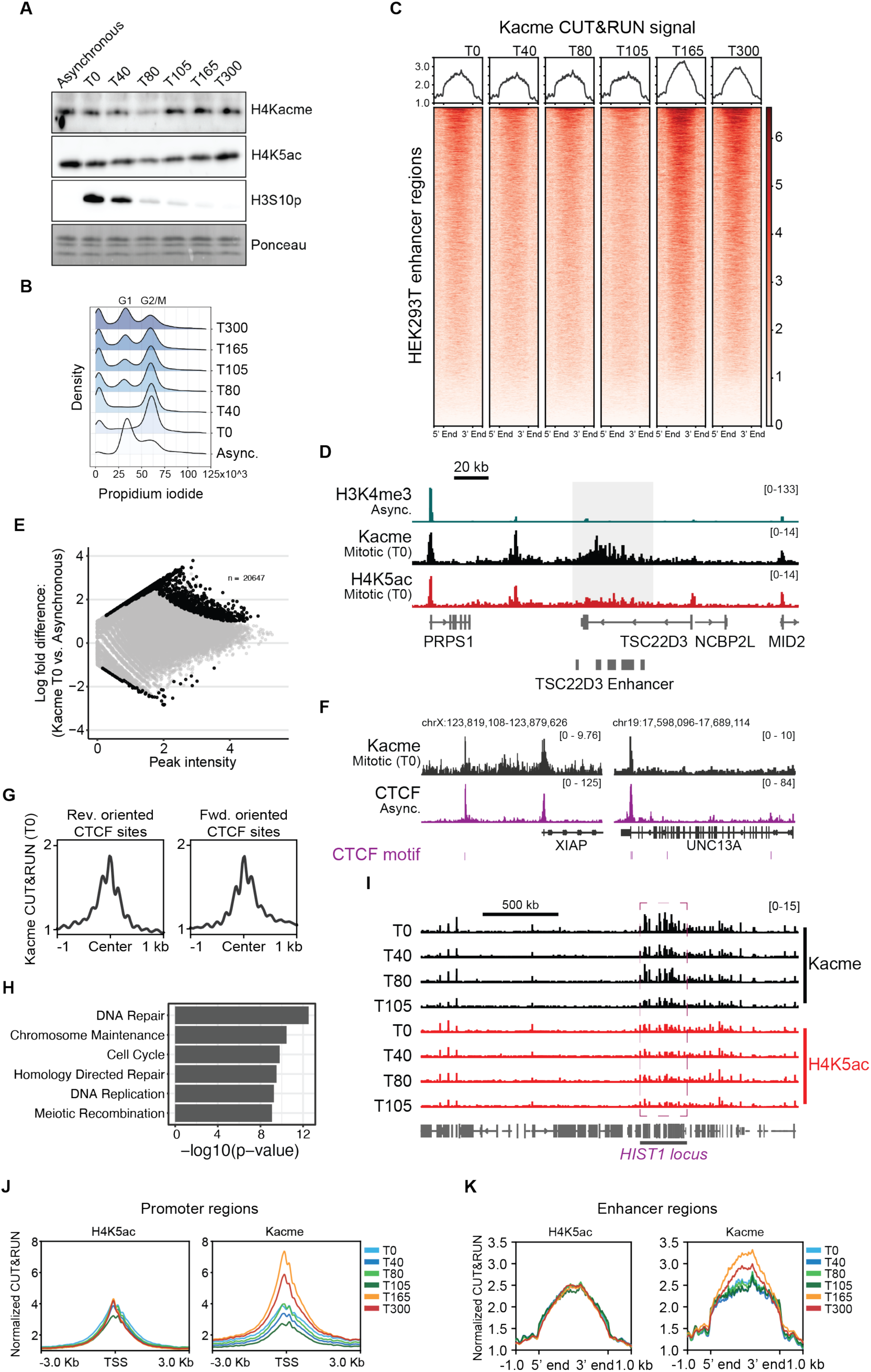
**(A)** Western blot analysis of H4Kacme, H4K5ac, and H3S10P in asynchronous HEK293T cells and HEK293T cells collected at various timepoints following release from nocodazole-mediated mitotic arrest. **(B)** Flow cytometry analysis of DNA content (propidium iodide) in asynchronous HEK293T cells and HEK293T cells collected at various timepoints following release from nocodazole-mediated mitotic arrest. **(C)** Heatmaps of normalized Kacme CUT&RUN signal at HEK293T-specific enhancer regions throughout the mitotic-release time course. **(D)** Genome browser tracks showing H3K4me3 in asynchronous cells and mitotic-Kacme/H4K5ac CUT&RUN signal at the TSC22D3 enhancer region (shaded in grey). **(E)** Differential binding analysis of regions of enrichment from Kacme CUT&RUN performed in mitotically arrested cells (T0) and asynchronous cells. Black points are regions of significant difference (|L2FC| > 1, p-value < 0.05). **(F)** Representative genome browser tracks showing CTCF enrichment in asynchronous HEK293T cells and Kacme CUT&RUN signal in mitotically arrested (T0) HEK293T cells. FIMO-predicted CTCF binding sites are annotated in magenta. **(G)** Aggregate Kacme CUT&RUN signal in mitotically arrested (T0) HEK293T cells at forward and reverse-oriented FIMO-predicted CTCF binding sites. **(H)** Biological pathway enrichment analysis for genes with significantly high relative promoter Kacme-to-H4K5ac ratios in mitotically arrested (T0) HEK293T cells. **(I)** Genome browser view of H4K5ac and Kacme CUT&RUN signal across the *HIST1* locus during the mitotic-release time course. **(J)** Metaplots of H4K5ac and Kacme CUT&RUN signal at promoter regions (±3 kb around TSS) across the mitotic-release time course. **(K)** Metaplots of H4K5ac and Kacme CUT&RUN signal at HEK293T-specific enhancer regions across the mitotic-release time course.

**Fig. S6.**
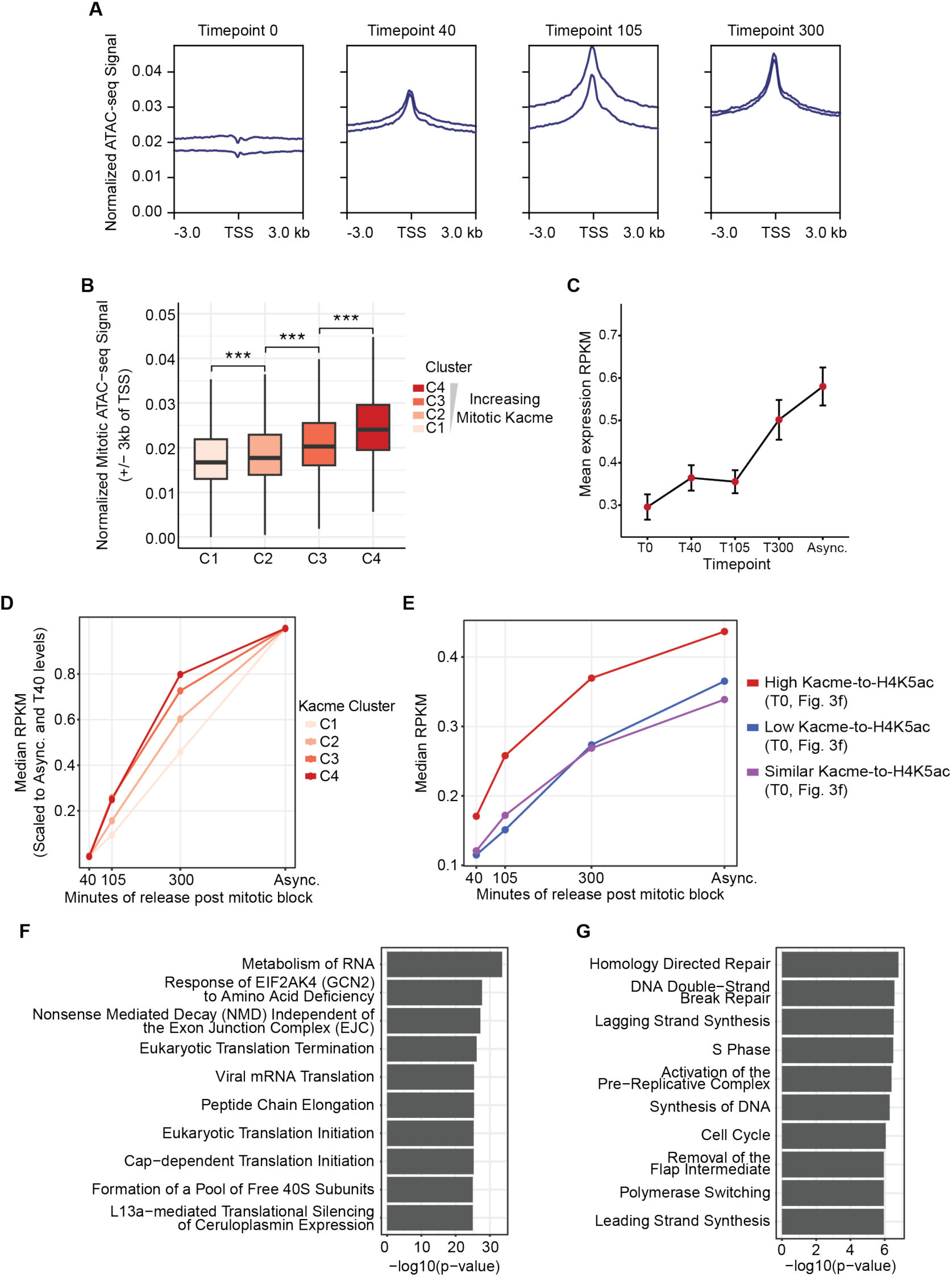
**(A)** Normalized ATAC-seq signal at four timepoints following mitotic release, centered on transcription start sites (±3 kb) across all annotated promoter regions. **(B)** Boxplots showing promoter ATAC-seq signal (±3 kb of TSS) in mitotically arrested (T0) HEK293T cells. Genes are grouped into four clusters (C1-C4) defined by mitotic Kacme levels (see Fig. 3D). Increasing Kacme is associated with higher chromatin accessibility in mitotically arrested cells. Statistical significance was determined using pairwise Wilcoxon rank-sum tests. *** = p < 0.001. **(C)** Mean nascent transcription (RPKM) across the mitotic-release time course, as assayed by TimeLapse-seq. Nascent transcription globally increases following mitotic exit, consistent with broad transcriptional reactivation. **(D)** Median RPKM values of nascent transcription for genes associated with each Kacme cluster (C1-C4; Fig. 3D) across mitotic release timepoints. Values were further scaled to reflect transcriptional reactivation relative to T40 and asynchronous levels, enabling direct comparison of reactivation dynamics across clusters. **(E)** Comparison of nascent transcriptional dynamics at genes preferentially marked by Kacme, H4K5ac, or both, during mitosis (T0). Differential binding analysis was used to identify promoters with significantly differential Kacme and H4K5ac enrichment in mitotically arrested cells (T0), as described in Fig. 3F. Median RPKM values of nascent transcription for genes associated with each set are shown across mitotic release timepoints. **(F)** Pathway enrichment analysis for genes with high relative mitotic Kacme-to-H4K5ac ratios (see Fig. 3J), that exhibit early reactivation following mitotic release. Genes were classified as early reactivation genes if they first reached nascent transcription levels ≥0.5-fold above asynchronous levels within 40 minutes of mitotic release. **(G)** Pathway enrichment analysis for genes with high relative mitotic Kacme-to-H4K5ac ratios (see Fig. 3J), that exhibit late reactivation following mitotic release. Genes were classified as late reactivation genes if they first reached nascent transcription levels ≥0.5-fold above asynchronous levels 300 minutes after mitotic release.

**Fig. S7.**
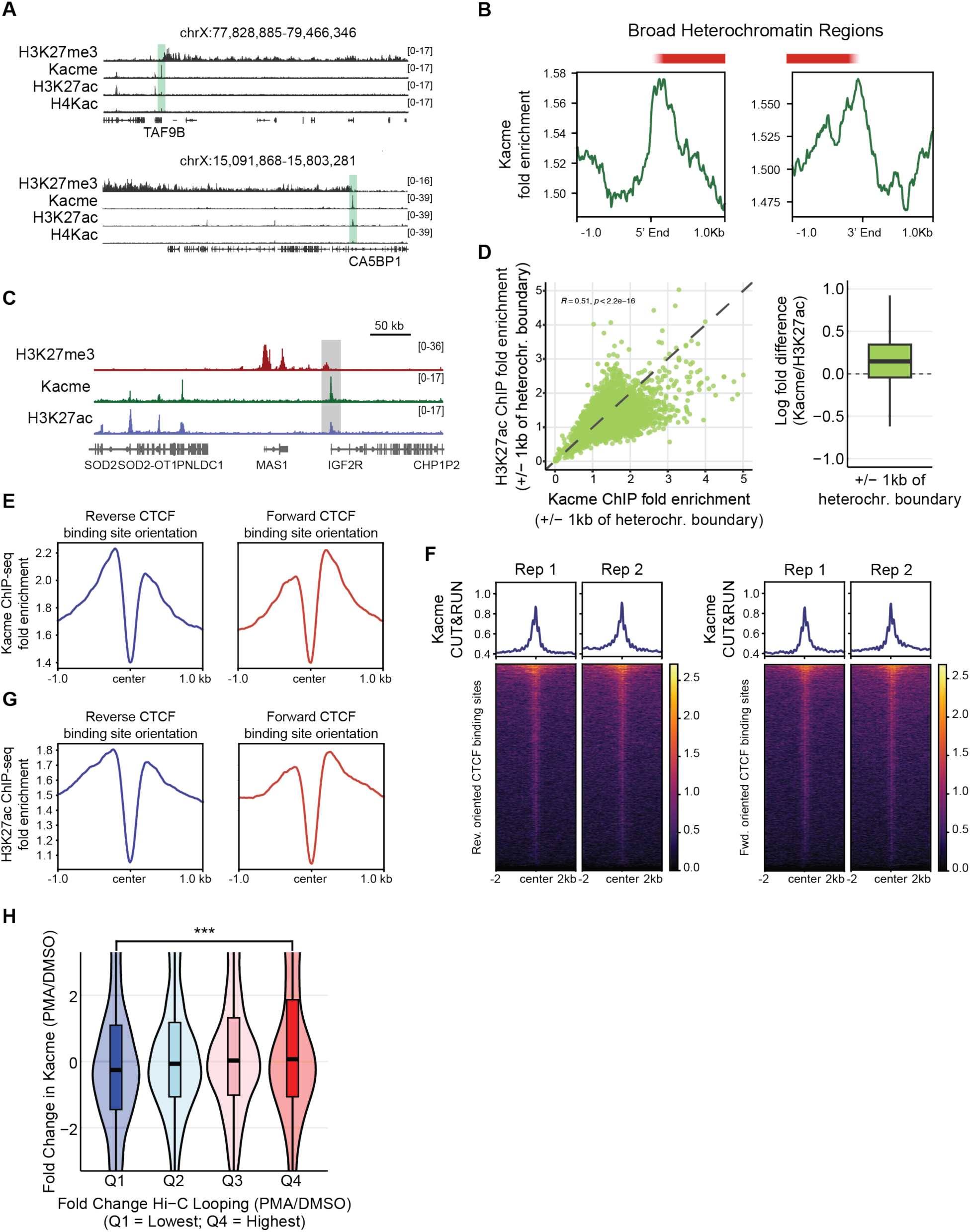
**(A)** Genome browser views showing HEK293T H3K27me3 CUT&RUN and Kacme, H3K27ac, and H4Kac ChIP-seq profiles at the borders of representative X chromosome heterochromatin domains. **(B)** Metagene analysis of HEK293T Kacme ChIP-seq enrichment at the 5’ and 3’ boundaries of broad H3K27me3 domains (n = 4,162). **(C)** Genome browser views showing H3K27me3 CUT&RUN and Kacme and H3K27ac ChIP-seq profiles at the border of a heterochromatin domain proximal to the *IGF2R* locus. **(D)** Left: Scatter plot comparing overall Kacme and H3K27ac ChIP-seq fold enrichment in HEK293T cells, measured within ±1 kb of n = 4,162 heterochromatin boundaries. Pearson correlation values and significance are indicated. Right: Boxplot showing the log fold difference in Kacme and H3K27ac enrichment at boundary-proximal regions. **(E)** Metaplots of Kacme ChIP-seq fold enrichment centered on forward- and reverse-oriented FIMO-predicted CTCF binding sites (± 1kb). **(F)** Heatmaps of Kacme CUT&RUN signal centered on forward- and reverse-oriented FIMO-predicted CTCF binding sites (± 2kb). CUT&RUN from two biological replicates are shown. **(G)** Metaplots of H3K27ac ChIP-seq fold enrichment centered on forward- and reverse-oriented FIMO-predicted CTCF binding sites (± 1kb). **(H)** Violin plots showing fold change in THP-1 promoter Kacme ChIP-seq signal (PMA/DMSO), binned by quartiles of changes in Hi-C loop strength (as reported in (*49*)). Genes exhibiting increased PMA-induced looping (Q4) are associated with increased Kacme signal, while genes exhibiting decreased looping (Q1) are associated with decreased Kacme. Statistical significance was determined using pairwise Wilcoxon rank-sum tests. *** = p < 0.001.

**Fig. S8.**
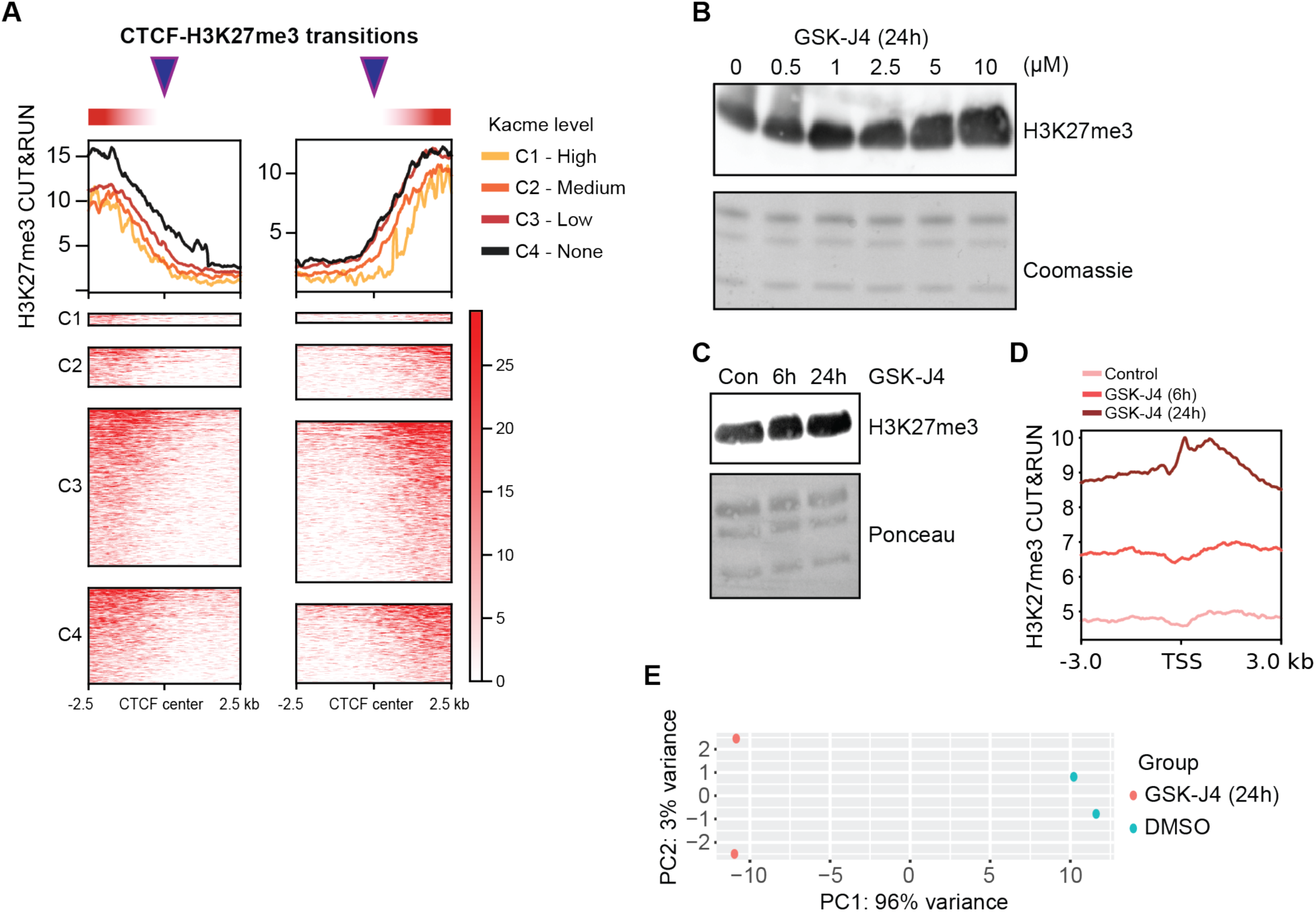
**(A)** Heatmaps showing H3K27me3 signal intensity in untreated HEK293T cells at CTCF binding sites that demarcate transitions in H3K27me3-enriched heterochromatin domains (L2FC H3K27me3 > 1). Transition sites were stratified by Kacme co-enrichment levels (C1-C4, see Fig. 6B). Transition sites with high Kacme levels (C1) exhibit reduced H3K27me3 spreading into neighboring euchromatin regions. **(B)** Representative western blot of global H3K27me3 levels following 24 h of GSK-J4 treatment (0-10 μM). **(C)** Representative western blot of global H3K27me3 levels following 6 or 24 h of GSK-J4 treatment (5 μM). **(D)** Metaplots of spike-in normalized H3K27me3 CUT&RUN signal in DMSO-treated HEK293T cells and HEK293T cells treated with 5 μM GSK-J4 for 6 h or 24 h. Inhibition of KDM6A/B results in a global increase in promoter H3K27me3 levels. **(E)** Principal component analysis (PCA) plot of spike-in normalized TimeLapse-seq read counts, transformed by the rlog function in DESeq2. DMSO- and 24 h GSK-J4-treated samples clustered separately.

**Table S1.**
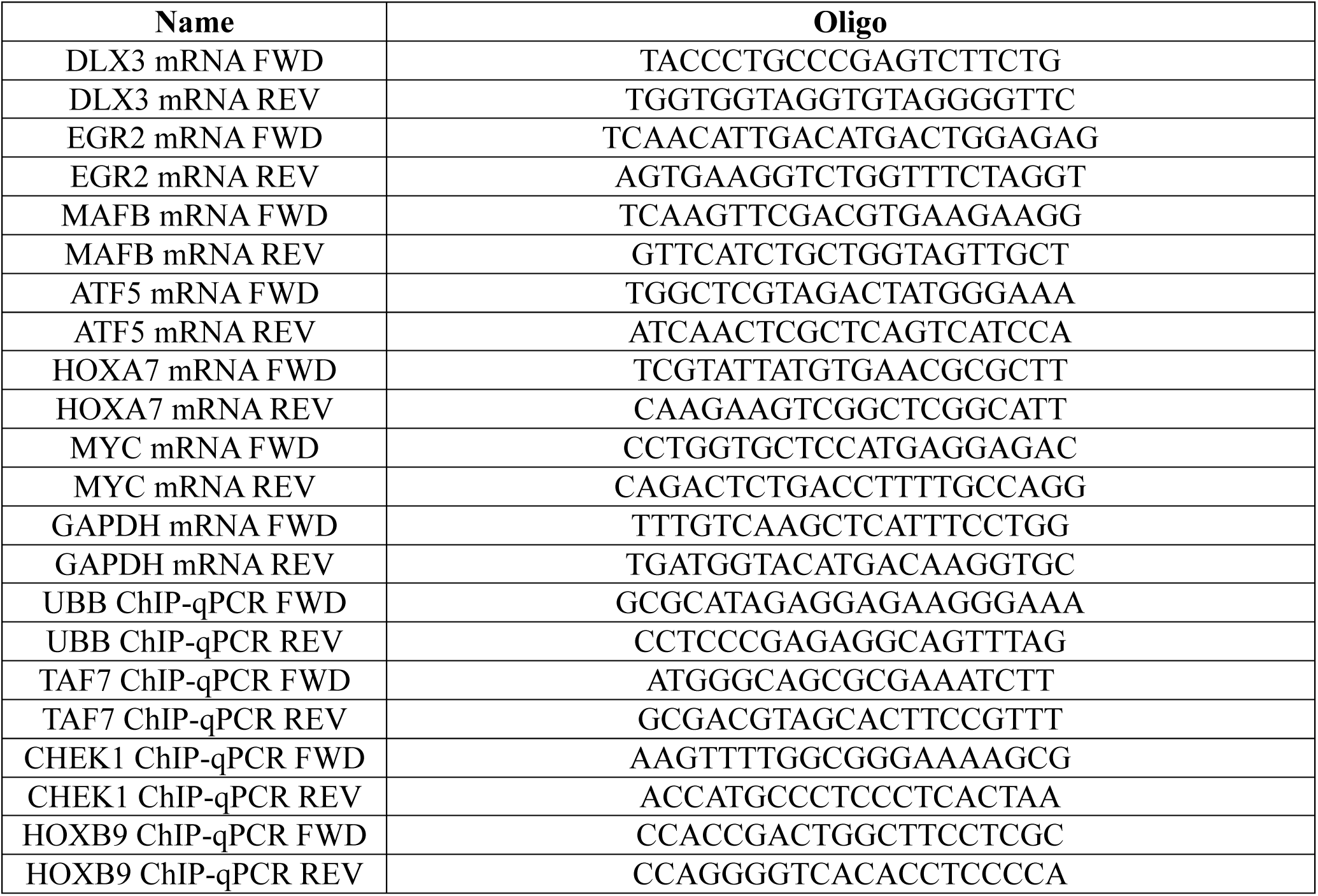
Oligos used in this study.

**Table S2.** Deposited data used in this study.

| **Resource** | **Source** | **Identifier** |
| --- | --- | --- |
| ATAC-seq in K562 | ENCODE | ENCSR868FGK |
| ATAC-seq in THP-1 monocyte; macrophage | Phanstiel et al., 2017 | GSE96800 |
| ATAC-seq in HEK293T | Pintado-Urbanc et al., 2026 | GSE313073 |
| ATAC-seq in S2 | Elrod et al., 2019 | GSE114467 |
| S2 H3K4me3, H3K4me1, H3K27ac ChIP-seq | Henriques et al., 2018 | GSE85191 |
| S2 Kacme ChIP-seq | Lu-Culligan et al., 2023 | GSE182204 |
| HEK293T Kacme, H4Kac ChIP-seq | Lu-Culligan et al., 2023 | GSE182204 |
| K562 Kacme ChIP-seq | Lu-Culligan et al., 2023 | GSE182204 |
| H3K27ac, CTCF ChIP-seq in THP-1 monocyte; macrophage | Phanstiel et al., 2017 | GSE96800 |
| K562 BRD2 ChIP-seq | ChIP-Atlas | SRX6913085 |
| H3K27me3, H3K9me3 ChIP-seq in THP-1 monocyte; macrophage | Lin et al., 2022 | GSE208046 |
| RNA-seq in THP-1 monocyte; macrophage | Lin et al., 2022 | GSE208046 |
| K562 Chromatin State Segmentation by HMM | ENCODE | ENCSR961HFL |
| H3K4me1 ChIP-seq in HEK293 | ENCODE | ENCSR000FCG |
| H3K4me3 ChIP-seq in K562 | ENCODE | ENCSR668LDD |
| H3K27ac ChIP-seq in K562 | ENCODE | ENCSR000AKP |
| H3K27me3 ChIP-seq in K562 | ENCODE | ENCSR000AKQ |
| H3K4me1 ChIP-seq in K562 | ENCODE | ENCSR000AKS |
| H3K36me3 ChIP-seq in K562 | ENCODE | ENCSR000AKR |
| H3K9me3 ChIP-seq in K562 | ENCODE | ENCSR000APE |
| H3K9ac ChIP-seq in K562 | ENCODE | ENCSR000AKV |
| H3K4me2 ChIP-seq in K562 | ENCODE | ENCSR000AKT |
| H2AZ ChIP-seq in K562 | ENCODE | ENCSR000APC |
| H3K79me2 ChIP-seq in K562 | ENCODE | ENCSR000APD |
| H4K20me1 ChIP-seq in K562 | ENCODE | ENCSR000AKX |
| CTCF ChIP-seq in K562 | ENCODE | ENCSR000AKO |
| POLR2A ChIP-seq in K562 | ENCODE | ENCSR388QZF |
| P300 ChIP-seq in K562 | ENCODE | ENCSR000EGE |
| RRBS in HEK293 | ENCODE | ENCSR794HFF |

